# Is disrupted sleep a risk factor for Alzheimer’s disease? Evidence from a two-sample Mendelian randomization analysis

**DOI:** 10.1101/609834

**Authors:** Emma L Anderson, Rebecca C Richmond, Samuel E Jones, Gibran Hemani, Kaitlin. H Wade, Hassan S Dashti, Jacqueline M Lane, Heming Wang, Richa Saxena, Ben Brumpton, Roxanna Korologou-Linden, Jonas B Nielson, Bjørn Olav Åsvold, Gonçalo Abecasis, Elizabeth Coulthard, Simon D. Kyle, Robin N Beaumont, Jessica Tyrrell, Timothy M Frayling, Marcus R Munafò, Andrew R Wood, Yoav Ben-Shlomo, Laura D Howe, Debbie A Lawlor, Michael N Weedon, George Davey Smith

**Affiliations:** MRC Integrative Epidemiology Unit at the University of Bristol, UK; Population Health Sciences, Bristol Medical School, University of Bristol, UK; Genetics of Complex Traits, University of Exeter Medical School, Exeter, UK; Center for Genomic Medicine, Massachusetts General Hospital, Harvard Medical School, Boston MA USA; Program in Medical and Population Genetics, Broad Institute, Cambridge MA USA; Department of Anesthesia, Critical Care and Pain Medicine, Massachusetts General Hospital, Boston, MA USA; Division of Sleep and Circadian Disorders, Brigham and Women’s Hospital, Harvard Medical School, Boston, MA, USA; K.G. Jebsen Center for Genetic Epidemiology, Department of Public Health and Nursing, Faculty of Medicine and health sciences, Norwegian University of Science and Technology, NTNU, Trondheim, Norway; Department of Thoracic Medicine, St. Olavs Hospital, Trondheim University Hospital, Trondheim, Norway; Department of Internal Medicine, Division of Cardiovascular Medicine, University of Michigan, Ann Arbor, MI, USA; Department of Endocrinology, St. Olavs Hospital, Trondheim University Hospital, Trondheim, Norway; Translational Health Sciences, Bristol Medical School, University of Bristol, UK; Sleep and Circadian Neuroscience Institute, Nuffield Department of Clinical Neurosciences, University of Oxford; UK Centre for Tobacco and Alcohol Studies, School of Psychological Science, University of Bristol, UK

## Abstract

**INTRODUCTION:** It is established that Alzheimer’s disease (AD) patients experience sleep disruption. However, it remains unknown whether disruption in the quantity, quality or timing of sleep is a risk factor for the onset of AD.

**METHODS:** Mendelian randomization (MR) was used to estimate the causal effect of self-reported and accelerometer-measured sleep parameters (chronotype, duration, fragmentation, insomnia, daytime napping and daytime sleepiness) on AD risk.

**RESULTS:** Overall, there was little evidence that sleep traits affect the risk of AD. There was some evidence to suggest that self-reported daytime napping was associated with lower AD risk (odds ratio [OR]: 0.70, 95% confidence interval [CI]: 0.50 to 0.99). Some other sleep traits (accelerometer-measured eveningness and sleep duration, and self-reported daytime sleepiness) had ORs for AD risk of a similar magnitude to daytime napping, but were less precisely estimated.

**DISCUSSON:** Our findings provide tentative evidence that daytime napping may reduce AD risk. However, findings should be replicated using independent samples.

## INTRODUCTION

Alzheimer’s disease (AD) has been estimated to affect 47 million people worldwide and the prevalence is expected to double in the next 20 years.[1] Current treatments are unable to reverse or delay progression of the disease, highlighting the importance of prevention. Identifying causal, modifiable risk factors is crucial for developing successful prevention strategies. It is well established that patients with AD experience sleep disruption (e.g. shorter duration, greater fragmentation[2–6]). However, it remains unknown whether disruption in the quantity, quality or timing of sleep is a causal risk factor for the onset of AD.

Various sleep parameters have been suggested as potential risk factors for AD in previous reviews and reports[7–10], but research to date has yielded inconsistent findings. Authors of the recent Lancet commission on ‘Dementia prevention, intervention, and care’, did not include sleep in their calculations of population attributable fractions of the most ‘potent’ dementia risk factors (despite acknowledging sleep as a potentially important risk factor) due to the absence of systematic reviews or enough consistent, high-quality evidence.[7] Inconsistencies in the sleep-AD literature may, at least in part, be explained by bias due to reverse causation. Most studies have been conducted in select clinical populations (e.g. patients with mild cognitive impairment or early AD) making it difficult to rule out that associations are not due to sleep disruption as a result of accumulating AD pathology. Few studies have been conducted in healthy (non-clinical) populations and, even those that have, tend to include older participants (i.e. mid-to late-life at baseline).[11, 12] AD has a long prodromal phase that can occur up to 20 years prior to diagnosis[13, 14]. Thus, even in apparently healthy populations, measuring participants’ sleep in later life makes it difficult to rule out the possibility that those participants with sleep disruption are those with prodromal AD. Another potential explanation for the inconsistencies may be the considerable heterogeneity in existing study designs, which have examined various exposures (e.g. sleep duration[12, 15, 16], time spent in sleep stages[12], fragmentation[17], insomnia[18] and frequency and duration of daytime napping[19] measured both subjectively and objectively) and outcomes (e.g. cognitive function at a single time-point[20, 21], cognitive decline over time[22], mild cognitive impairment[23], AD diagnoses[12, 23] and putative AD biomarkers such as amyloid-beta and tau[17, 19]). Finally, all studies conducted to date are observational, meaning confounding is a plausible explanation for the findings.

Mendelian randomization (MR) is a method that uses genetic variants as instrumental variables for environmental modifiable exposures.[24] Due to their random allocation at conception, genetic markers of a risk factor are largely independent of potential confounders that may otherwise bias the association of interest.[25] They also cannot be modified by subsequent disease, thereby eliminating potential bias by reverse causation. Thus, MR is a useful tool for helping to establish whether sleep traits are causally related to risk of future AD, or whether associations observed to date are likely to be a result of bias by confounding and/or reverse causation. In this study, we aimed to establish whether both self-report and accelerometer-measured sleep traits have a causal effect on AD risk, using a two-sample MR design[26].

## METHODS

Methods for conducting two-sample MR analyses have been published previously[26]. Briefly, two-sample MR provides an estimate of the causal effect of an exposure on an outcome, using independent samples to obtain the gene-exposure and gene-outcome associations, provided three key assumptions hold: (i) genetic variants are robustly associated with the exposure of interest (i.e. replicate in independent samples), (ii) genetic variants are not associated with potential confounders of the association between the exposure and the outcome and (iii) there are no effects of the genetic variants on the outcome, independent of the exposure (i.e. no horizontal pleiotropy)[27].

### Samples

Genome-wide association studies (GWAS) were previously performed for seven self-reported measures of habitual sleep patterns including chronotype[28], sleep duration[29], long sleep duration[29], short sleep duration[29], frequent insomnia[30], excessive daytime sleepiness and daytime napping. GWAS have also been performed for three accelerometer-measured measures of sleep including timing of the least active 5 hours of the day (L5 timing), nocturnal sleep duration and sleep fragmentation[31]. Note that the assessment of accelerometer-derived sleep for up to 7 days per individual in UK Biobank was performed, on average, five years after the self-report sleep data were collected.[32] Table 1 provides (i) a description of each of the sleep traits and the units in which they were measured, (ii) participant numbers for each GWAS, (iii) the number of approximately independent genome-wide significant (p<5×10^−8^) loci identified by each GWAS and (iv) the F statistic for each trait. F statistics provide an indication of instrument strength[33] and are a function of R^2^ (how much variance in the trait is explained by the set of genetic instruments being used), the number of instruments being used, and the sample size. F>10 is typically used as a threshold to indicate that weak instrument bias will have a small influence on the causal effect estimate[34]. For the outcome, we used the large-scale GWAS meta-analysis of AD, conducted by the International Genomics of Alzheimer’s Project (IGAP) (n=17,008 AD cases and 37,154 controls).[35] Ethics approval was obtained by the original GWAS studies.

**Table 1:** Description of the sleep GWAS included in the two-sample MR analyses.

|  | **Trait definition (units)** | **N Participants**<br>**(cases/controls**<br>**for binary traits)** | **N loci**<br>**identified in**<br>**GWAS** | **F**<br>**Statistic** |
| --- | --- | --- | --- | --- |
| **Self-reported measures** |  |  |  |  |
| **Chronotype[28]** | Whether a person identifies as being a 'morning person' or an 'evening person' (ordered categorical variable of definitely a morning person, more a morning than an evening person, do not know, more an evening than morning person and definitely an evening person)* | 449,734 | 351 | 33.1 |
| **Sleep duration[29]** | Average number of hours slept in 24 hours, including naps (continuous variable, hours) | 446,118 | 78 | 39.6 |
| **Short sleep duration[29]** | Person has an average of 6 hours or less per night vs 7-8 hours per 24 hours (binary variable of yes/no) | 411,934<br>(106,192/305,742) | 27 | 25.6 |
| **Long sleep duration[29]** | Person has an average of 9 hours or more per night vs 7-8 hours per 24 hours (binary variable of yes/no) | 339,926<br>(34,184/305,742) | 8 | 30.6 |
| **Frequent insomnia[30]** | Person has trouble falling asleep at night or wakes up in the middle of the night (binary variable of usually vs never/rarely) | 453,379<br>(131,480/321,899) | 48 | 41.7 |
| **Excessive daytime sleepiness** | Person dozes off or falls asleep during the day without meaning to (ordered categorical variable of never or rare, sometimes, often and all the time) | 452,071 | 37 | 42.3 |
| **Daytime napping** | Person naps during the day (ordered categorical variable of never, sometimes, and usually) | 452,633 | 170 | 46.3 |
| **Accelerometer measures** |  |  |  |  |
| **L5 timing[31]** | Timing of the least active 5 hours of the day (continuous variable of hours elapsed since previous midnight; provides indication of phase of most restful hours with later times indexing greater tendency towards 'eveningness') | 85,205 | 6 | 55.3 |
| **Sleep duration[31]** | Average number of hours of nocturnal sleep per night (continuous variable, hours) | 84,810 | 11 | 52.1 |
| **Sleep fragmentation[31]** | The average number of nocturnal sleep episodes separated by at least 5 minutes of wakefulness per night (continuous variable, number of episodes) | 84,810 | 21 | 38.6 |

\*Note that in the original chronotype GWAS, categories were ordered from more 'eveningness' to more 'morningness'. In this analysis, to ensure that the ordinal chronotype variable correlated positively with the accelerometer-measured measure of L5 timing, SNP-exposure coefficients for chronotype were reordered from more 'morningness' to more 'eveningness' (where 'definitely a morning person' is the reference category).

## STATISTICAL ANALYSIS

### Harmonization of exposure and outcome GWAS data

Only biallelic single nucleotide polymorphisms (SNPs) were included as instruments (insertions and deletions were excluded). SNPs for each sleep trait were identified in the AD GWAS dataset. Proxies were identified for any SNPs not found (r^2^>0.8 using 1000 genomes as a reference). In a two-sample MR analysis, the effect of a SNP on exposure and an outcome must be harmonised relative to the same allele. SNPs for the exposure were coded so that the effect allele was always the ‘increasing allele’ (e.g. increasing sleep duration), and the alleles were harmonized so that the effect on the outcome corresponded to the same allele as the exposure. Supplemental Table A shows the SNP flow through the harmonization procedure and the final number of SNPs included in each analysis. Phenotypic correlations between each of the sleep traits were estimated using Pearson’s *r*.

### Estimating the causal effects of the sleep phenotypes on risk of Alzheimer’s disease

MR-Base (www.mrbase.org)[36] was employed to perform all two-sample MR analyses. Effect estimates and corresponding standard errors of the genome-wide significant SNPs were extracted from each sleep GWAS and the AD GWAS. The SNP-exposure (sleep trait units detailed in Table 1) and SNP-outcome (AD, in units of log odds ratios [ORs]) coefficients were combined using an inverse-variance-weighted (IVW) approach to give an overall estimate of the causal effect across all SNPs included for each sleep phenotype. This is equivalent to a weighted regression of the SNP-outcome coefficients on the SNP-exposure coefficients with the intercept constrained to zero. The results of all analyses were converted to ORs for AD. For binary exposures (i.e. frequent insomnia, and long and short sleep duration), SNP-exposure coefficients were estimated using logistic regression and are therefore on the log odds scale. Causal effect estimates (i.e. ORs for AD) have been rescaled so that they are interpreted per doubling of genetic liability for the sleep trait, as recommended by Burgess et al[37]. For ordered categorical exposures, SNP-exposure coefficients were estimated using linear regression and causal effect estimates are interpreted per category increase in the sleep trait. For continuous exposures, SNP-exposure coefficients were estimated using linear regression and causal effect estimates are interpreted per unit increase in the sleep trait (units detailed in Table 1).

### Sensitivity analyses

A series of sensitivity analyses were conducted to check for violation of the key MR assumptions. The rationale and methodological details for each of these analyses are provided in the online supplement. The IVW method assumes no horizontal pleiotropy but will be unbiased if there is balanced horizontal pleiotropy.[27] Thus, results from the IVW method were compared to those from MR-Egger[27] and weighted median regressions[38] which relax this assumption. The IVW method also assumes no measurement error in the gene-exposure association estimates (i.e. the NOME assumption)[27]. We assessed this using an adaptation of the I^2^ statistic[39] (referred to as I^2^_GX_), which provides an estimate of the degree of regression dilution in the MR-Egger causal estimate due to uncertainty in the SNP-exposure estimates. Simulation extrapolation (SIMEX) was then used to adjust the MR-Egger estimate for this dilution[40]. Heterogeneity (i.e. variability in causal estimates from different genetic variants) was assessed using Cochran’s Q statistic[27]. Funnel plots were then generated to enable visual assessment of the extent to which pleiotropy is likely to balanced (or directional) across the set of instruments used in each analysis. Radial MR was used to detect and remove any SNP outliers (i.e. those SNPs that contribute the most heterogeneity to Cochran’s Q, based on a multiple testing corrected P value threshold). Leave-one-out permutations were conducted to assess the undue influence of potentially pleiotropic SNPs on the causal estimates[41]. We checked results were similar after excluding palindromic SNPs[42]. Steiger filtering was performed to test that the hypothesised causal direction was correct for each SNP (i.e. that that the genetic instruments influence the exposure first and then the outcome, through the exposure)[43]. Finally, we investigated potential bias due to ‘winner’s curse’; where the magnitude of the effect sizes for variants identified within a single discovery sample are likely to be larger than in the overall population, even if they are truly associated with the exposure. Assessment of winner’s curse was only possible for frequent insomnia and sleep duration, where the GWAS were replicated in independent samples (The Nord-Trøndelag Health Study [HUNT] and The Cohorts for Heart and Aging Research in Genomic Epidemiology [CHARGE], respectively). Details of the replication samples and winners curse analyses are in the online supplement.

### Additional analyses

Causal effects of the of sleep traits on risk of AD were additionally assessed by two-sample Mendelian randomization analysis using summary statistics from the most recently published AD GWAS meta-analysis[44]. This includes the aforementioned IGAP, the AD working group of the Psychiatric Genomics Consortium and the AD Sequencing Project (Phase 1; n= 24,087 cases and 55,058 controls compared with n=17,008 AD cases and 37,154 controls for IGAP alone). The paper details a further meta-analysis additionally including participants from the UK Biobank (Phase 3). However, we a-priori decided to use the Phase 1 rather than Phase 3 meta-analysis for two reasons: Firstly, all sleep trait GWASs include the UK Biobank, meaning there would be significant overlap between the exposure and outcome samples in each MR analysis (which can yield biased causal effect estimates[45]). Secondly, the phase 3 AD GWAS meta-analysis includes only AD-by-proxy cases from the UK Biobank (i.e. no diagnosed cases). AD-by-proxy cases were defined as a positive response to the question ‘Has your mother or father ever suffered from Alzheimer’s disease/Dementia’. There are several potential problems with this for MR analyses: participants defined as cases have not themselves been diagnosed with AD; the question does not specify Alzheimer’s disease but asks about any form of dementia; and finally, the question does not ask if family members were diagnosed by a doctor.

To examine whether the previously observed associations between AD (as an exposure) and sleep disruption (as an outcome)[2–6] could be supported through applying MR analytical approaches, we tested whether genetic liability for AD was causally associated with the self-reported and accelerometer-measured sleep traits, using 20 independent genome-wide significant AD SNPs identified in the IGAP AD GWAS meta-analysis (described previously)[35]. As with the main analysis of sleep traits on AD risk, SNP-exposure and SNP-outcome coefficients were combined using an inverse-variance-weighted (IVW). MR-Egger, weighted median, Radial MR and Steiger filtering were performed to assess potential violation of the MR assumptions. Analyses were conducted both with and without the ApoE variant included, as ApoE has been previously shown to be pleiotropic[46] (which violates an MR assumption). We also examined associations of ApoE (as a single genetic instrument) with the sleep traits. As AD is a binary exposure and SNP-exposure coefficients are on the log odds scale, causal estimates for the effect of AD on sleep traits are rescaled so that they are interpreted per doubling of genetic liability for AD. It is worth noting that there are several important limitations to these analyses (including the healthy selected, relatively young population in the UK Biobank) and these are discussed in detail in the online supplement. We present these results for completeness.

## RESULTS

Supplemental Table B shows the phenotypic correlations between each of the sleep traits. Correlations between accelerometer measures have been published previously.[31] Correlations were generally weak, ranging from r=-0.001 (between accelerometer-measured L5 timing and accelerometer-measured sleep duration) to r=-0.32 (between self-report frequent insomnia and self-report short sleep duration). It is also worth noting that correlations were weak between self-reported and accelerometer-measured sleep duration (*r*=0.15). This may reflect that accelerometer data in the UK biobank were collected between 2 and 9 years (mean 5 years)[31] after baseline, when self-reported sleep measures were assessed. It may also reflect self-reports of global sleep duration (vs daily self-reported) can be influenced by distress/affect[47].

Figure 1 shows results for the IVW analysis of the sleep traits on risk of AD. Full results can be found in Supplemental Table C. Point estimates for self-reported chronotype, insomnia and sleep duration were very close to or on the null. Both shorter and longer self-reported sleep duration and accelerometer-measured sleep fragmentation yielded positive estimates with AD risk, but were imprecisely estimated. Similarly, the estimates suggesting protective effects for self-reported daytime napping was associated with lower AD risk, with odds of AD being 30% lower [95% CI: 1% to 50%] per category increase from ‘never’, ‘sometimes’ to ‘usually’ napping, were also imprecisely estimated. Odds ratios for greater accelerometer-measured eveningness, longer accelerometer-measured sleep duration and self-reported daytime sleepiness were similar in magnitude to daytime napping, although again with wide confidence intervals.

**Figure 1:**
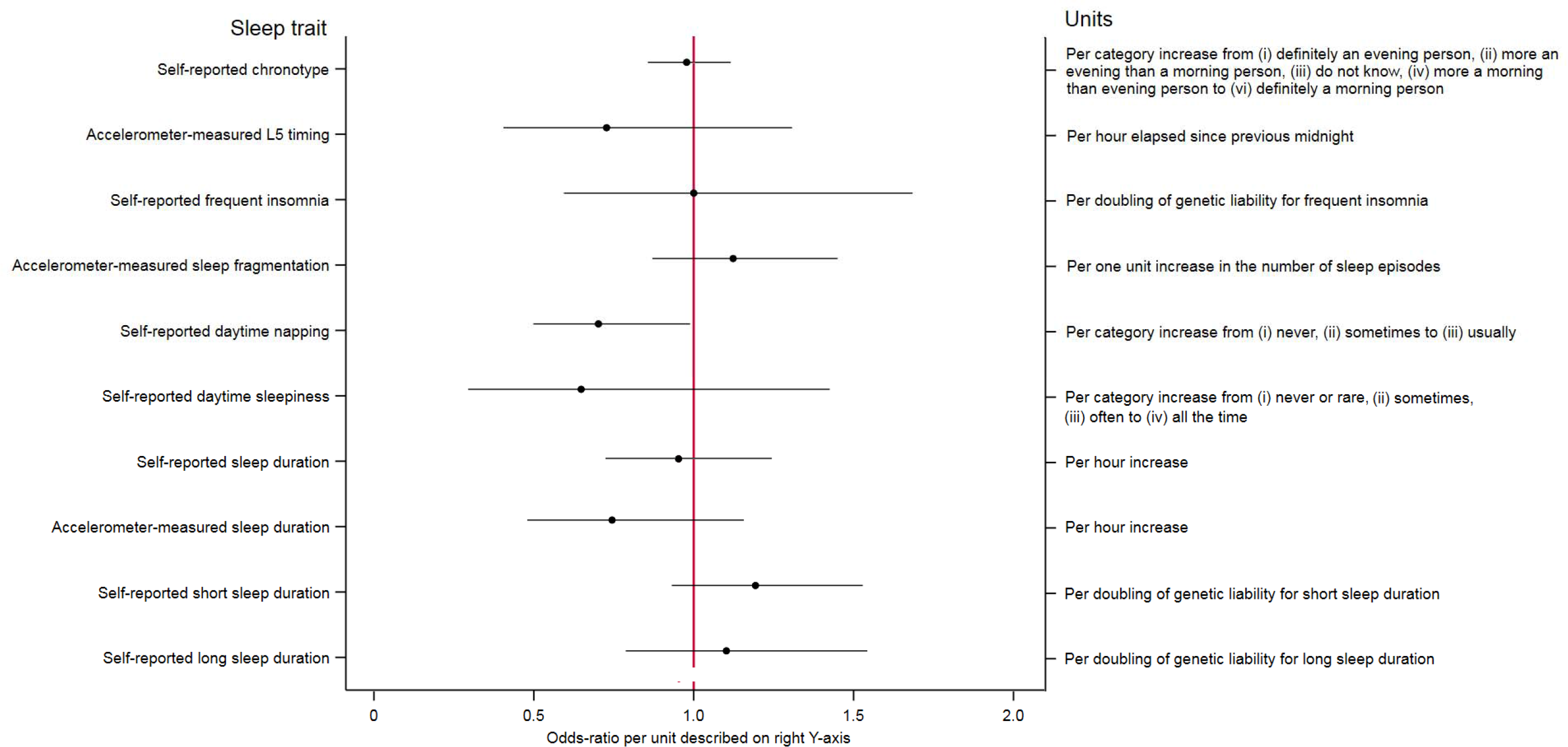
Associations of sleep traits with Alzheimer’s Disease.

### Sensitivity analyses

For all analyses, there was little evidence of directional pleiotropy from the MR-Egger regression intercepts (Supplemental Table D) and causal effect estimates from the MR-Egger and weighted median regressions generally agreed with those from the IVW regressions; in all cases there was substantial overlap between the confidence intervals for each estimate (Supplemental Table C). As expected, precision was less for MR-Egger (due to estimating both an intercept and slope in the MR-Egger regression as opposed to only a slope in the IVW regression) and weighted median (due to assuming only 50% of the instruments are valid). I^2^_GX_ statistics are provided in Supplemental Table E and SIMEX-adjusted MR-Egger estimates in Supplemental Table C. These estimates were consistent with regression dilution of the MR-Egger causal effect estimates due to measurement error in the SNP-exposure estimates. There was evidence of between-SNP heterogeneity in the self-reported chronotype and daytime napping, and accelerometer-measured L5-timing analyses (Supplemental Table F). However, there were not unduly asymmetrical in the funnel plots (Supplemental Figures A-C), suggesting directional pleiotropy is unlikely to bias the effect estimates for these sleep traits. A total of three outliers were detected by Radial MR for sleep fragmentation (rs12714404, rs429358 and rs4974697) and one for L5 timing (rs1144566). Point estimates for these two traits attenuated towards the null after removal of these outliers (Supplemental Table G). Results were similar after removing each SNP in turn in the leave-one-out permutations (Supplemental Figures D-M), suggesting no single SNP was having undue influence on the overall causal effect estimate. Results were also similar when palindromic SNPs were excluded from the analyses (Supplemental Table H). Steiger filtering provided evidence that for each MR analysis, SNPs explained more variation in the sleep trait than in AD. Findings for the sleep duration and insomnia results were similar when repeating analyses using the available replication datasets (i.e. using SNP-exposure estimates from independent datasets) (Supplemental Table I), providing evidence that bias due to winner’s curse is unlikely.

### Additional analyses

Causal effect estimates for associations of sleep traits on risk of AD were very similar when using the largest meta-analysis GWAS for AD, typically with more precision around the causal estimates (Supplementary Figure D). There was consistent evidence of a protective effect of daytime napping on Alzheimer’s risk, with odds of AD being 36% lower [95% CI: 11% to 45%] per category increase from ‘never’, ‘sometimes’ to ‘usually’ napping in the IVW analysis. Effects were consistent across a number of pleiotropy-robust methods, including MR Egger and weighted median approaches.

Associations between genetic liability for AD and all sleep traits are provided in Supplemental Table J. All point estimates are interpreted per doubling of genetic risk for AD. There was little evidence that genetic liability for AD was associated with short sleep duration (<6 hours vs 7-8 hours per 24 hours) or daytime sleepiness. Increased genetic liability for AD was associated with less frequent insomnia, reduced daytime napping and reduced sleep fragmentation. The effects were very small in magnitude (e.g. a doubling of genetic liability for AD was associated with, on average, a 0.4% lower risk of frequent insomnia). Point estimates for associations of genetic liability for AD with all other sleep traits were also very close to or on the null (e.g. a doubling of genetic liability for AD was associated with, on average, 0.3 minutes lower sleep duration). P values were small; likely due to the strength of ApoE as an instrument for AD. Given that these causal effect estimates are per doubling of genetic liability for AD, the magnitude of effect is very small and not likely to be clinically important. Results were similar when using ApoE alone as an instrument for AD (i.e. excluding all other AD SNPs, Supplemental Table K). When using the full set of AD genetic instruments minus ApoE, point estimates remained largely unchanged but confidence intervals were wider (Supplemental Table L). Results were comparable when using MR-Egger and weighted median regressions (Supplemental Table J) and after removal of outliers detected by Radial MR (Supplemental Table M). Steiger filtering provided evidence that for each MR analysis, all AD SNPs explained more variation in AD than in the sleep trait.

Given our concerns about selection bias in the UK Biobank for these analyses[48], we performed a post-hoc analysis to assess whether causal effect estimates were comparable when using a different outcome sample. Methods for these analyses are provided in the online supplement. We tested the association between genetic liability for AD and frequent insomnia in N=62,533 participants from The Nord-Trøndelag Health Study (HUNT)[30, 49]. HUNT is a less selected sample with over 60% response rate (compared with <5% for UK Biobank). Results were comparable to the main analyses using UK Biobank, except that confidence intervals were wider (IVW odds ratio: 0.98 per doubling of genetic liability for AD, 95% CI: 0.95 to 1.01 in HUNT vs IVW odds ratio: 0.99 per doubling of genetic liability for AD, 95% CI: 0.99 to 1.00 in the UK Biobank, Supplemental Table N).

## DISCUSSION

We have used the largest genome-wide association studies available of self-report and accelerometer-measured sleep traits and diagnosed Alzheimer’s disease, to provide bounds on possible causal relationships between them. Overall, based on the data presented in this study, we found no clear evidence of a link between these sleep traits and risk of AD, although there was suggestive evidence that daytime napping was associated with lower risk of AD. Odds ratios for accelerometer-measured ‘eveningness’, accelerometer-measured sleep duration, and self-reported daytime sleepiness were similar in magnitude to daytime napping, however, confidence intervals were wide and consistent with the null.

We found increased genetic liability for AD was associated with less frequent insomnia, reduced daytime napping and reduced sleep fragmentation. However, it is important to note that point estimates were very close to the null, and given that they are per doubling of genetic liability for AD, are unlikely to be clinically important. Confidence intervals around these point estimates were likely narrow due to the inclusion of ApoE as a genetic instrument for AD (as it is strongly and robustly associated with a 3- to 15-fold increase in AD risk[44]); confidence intervals were wider when we excluded this instrument. Although these associations are in the opposite direction to what we expected, these results should be treated with caution because of the potential for selection bias within the UK Biobank (which can cause spurious associations[48]), where participants are healthier than the general population. We observed a very small reduction in insomnia related to AD liability in an independent sample (HUNT), though with wider confidence intervals given its smaller size.

It is worth noting that we looked only at frequency (not duration) of daytime napping, as information on duration of naps is not currently available in the UK Biobank. Previous studies have reported both positive and negative outcomes observed in relation to napping, and there is evidence that the duration may be particularly important (with shorter naps being beneficial, and longer naps being detrimental for various health outcomes including cardiovascular risk, cognitive impairment and memory consolidation).[50–53] Most studies to date linking daytime napping to poor health outcomes have been done in elderly populations[4, 54, 55], making it difficult to rule out that (in those studies) daytime napping is not merely a result of underlying disease, rather than being a cause of the poor outcomes. In addition, it’s often not possible in those studies to separate whether it is poor nocturnal sleep or the consequential daytime napping that is associated with the adverse outcomes. Conversely, there is some evidence that daytime naps offer a variety of benefits including memory consolidation[56–58], improvements in subsequent learning[59], executive functioning[60] and emotional processing[61], all of which are impaired in AD[62, 63]. There is also evidence that short, restorative naps (<60 minutes in duration) may reduce the rate of cardiovascular disease and low-grade inflammation.[53] Thus, daytime napping (prior to the onset of preclinical disease) may potentially serve as a useful compensatory mechanism for poor nocturnal sleep, enabling the brain to carry out tasks it was unable to complete during poor nocturnal sleep (for example, due to fragmented or short sleep).

### Strengths and Limitations

To our knowledge, this is the first study to examine the causal effect of various sleep traits on risk of AD, using MR. We have both self-reported and accelerometer-assessed measures of sleep, allowing a comprehensive evaluation of various sleep parameters and a comparison across the two methods of assessment. We conducted a comprehensive series of sensitivity analyses to examine whether our results were robust to the various assumptions of MR or were likely to be biased by horizontal pleiotropy. We were also able to replicate some of our findings using independent exposure datasets (i.e. for self-report sleep duration and insomnia), and results were consistent, suggesting our findings are unlikely to be biased by winner’s curse. Findings were also largely consistent when we performed MR using summary statistics from a more recent GWAS meta-analysis of Alzheimer’s disease[44]. There are, however, several limitations to our study. The first limitation is low instrument strength; although F statistics were over 10 (the threshold typically used to indicate potential weak instrument bias), confidence intervals for analyses of sleep traits on risk of AD were generally wide. Identifying stronger instruments for these sleep traits may enable us to estimate their causal effects on AD risk more precisely. Secondly, there was a small number of approximately independent genome wide significant SNPs available for the MR analyses for some sleep traits (e.g. self-report long-sleep duration and for accelerometer L5 timing and sleep duration), making it difficult to examine potential directional horizontal pleiotropy using funnel plots. However, given that there was only evidence of heterogeneity for L5 timing, daytime napping and chronotype (and the chronotype and daytime napping showed no marked asymmetry in the funnel plot), horizontal pleiotropy is unlikely to explain our findings. Thirdly, we did not correct for multiple testing because several of the sleep traits are correlated, but the results need to be interpreted in this light and replication of our findings are required. Fourthly, it is possible that the specific features of sleep that are implicated in the pathogenesis of AD (for example, disruption of slow wave sleep) are not detectable using accelerometers or subjectively. Fifth, there may be a threshold effect of daytime napping by which shorter naps are beneficial and long, frequent naps may be detrimental. Previous studies have suggested this may be the case (particularly for cardiovascular risk, cognitive impairment, and memory consolidation[53]), but we are unable to unpick these effects with current data in an MR framework. Sixth, most analyses were conducted using data from the UK Biobank, which may not be representative of the general population (due to selection into the study)[64]. That said, results were similar when using data from independent replication samples (including HUNT and CHARGE). Finally, no sleep diaries were collected in the UK Biobank to identify time in bed and out of bed, which may introduce measurement error into some of the accelerometer measures. However, for the sleep data used in this study, time in and out of bed were estimated using a validated algorithm to determine the sleep period time window[65].

### Conclusions

Our findings tentatively suggest that daytime napping may reduce AD risk. However, given that this is the first MR study of multiple self-report and objective sleep traits on AD risk, findings should be replicated using independent samples. Identifying stronger instruments for all sleep traits will be useful in more precisely estimating any causal effects on AD risk.

## Author contributions

ELA conceptualised the study. ELA and RCR completed all statistical analyses with guidance from GDS, DAL, MNW, GH, LDH, KHW, SEJ and RCR. ELA drafted the first version of the manuscript. All authors provided critical comments on the manuscript. ELA is the guarantor and accepts full responsibility for the work and/or the conduct of the study, had access to the data, and controlled the decision to publish. The corresponding author attests that all listed authors meet authorship criteria and that no others meeting the criteria have been omitted.

## Acknowledgements

This research has been conducted using the UK Biobank Resource.

## References

[1] Prince M, Wilmo A, Guerchat M, Ali G, Wu Y, Prina M. World Alzheimer Report 2015. The Global Impact of Dementia: An analysis of prevalence, incidence, cost and trends. London2015. p. 1–87.

[2] Liguori C, Nuccetelli M, Izzi F, Sancesario G, Romigi A, Martorana A, et al. Rapid eye movement sleep disruption and sleep fragmentation are associated with increased orexin-A cerebrospinal-fluid levels in mild cognitive impairment due to Alzheimer’s disease. Neurobiol Aging. 2016;40:120–6.

[3] Beaulieu-Bonneau S, Hudon C. Sleep disturbances in older adults with mild cognitive impairment. Int Psychogeriatr. 2009;21:654–66.

[4] Cross N, Terpening Z, Rogers NL, Duffy SL, Hickie IB, Lewis SJ, et al. Napping in older people ‘at risk’ of dementia: relationships with depression, cognition, medical burden and sleep quality. J Sleep Res. 2015;24:494–502.

[5] da Silva RA. Sleep disturbances and mild cognitive impairment: A review. Sleep Sci. 2015;8:36–41.

[6] Guarnieri B, Adorni F, Musicco M, Appollonio I, Bonanni E, Caffarra P, et al. Prevalence of sleep disturbances in mild cognitive impairment and dementing disorders: a multicenter Italian clinical cross-sectional study on 431 patients. Dement Geriatr Cogn Disord. 2012;33:50–8.

[7] Livingston G, Sommerlad A, Orgeta V, Costafreda SG, Huntley J, Ames D, et al. Dementia prevention, intervention, and care. Lancet. 2017;390:2673–734.

[8] Prince M, Albanese E, Guerchet M, Prina M. World Alzheimer’s Report 2014. Dementia and Risk Reduction: An Analysis of Protective and Modifiable Factors. 2014.

[9] Baumgart M, Snyder HM, Carrillo MC, Fazio S, Kim H, Johns H. Summary of the evidence on modifiable risk factors for cognitive decline and dementia: A population-based perspective. Alzheimers Dement. 2015;11:718–26.

[10] Reed S, Wittenberg R, Karagiannidou M, Anderson R, Knapp M. Public Health England Report: The effect of midlife risk factors on dementia in older age. 2017.

[11] Ferrie JE, Shipley MJ, Akbaraly TN, Marmot MG, Kivimaki M, Singh-Manoux A. Change in sleep duration and cognitive function: findings from the Whitehall II Study. Sleep. 2011;34:565–73.

[12] Pase MP, Himali JJ, Grima NA, Beiser AS, Satizabal CL, Aparicio HJ, et al. Sleep architecture and the risk of incident dementia in the community. Neurology. 2017;89:1244–50.

[13] Rodriguez-Vieitez E, Saint-Aubert L, Carter SF, Almkvist O, Farid K, Scholl M, et al. Diverging longitudinal changes in astrocytosis and amyloid PET in autosomal dominant Alzheimer’s disease. Brain. 2016;139:922–36.

[14] Bateman RJ, Xiong C, Benzinger TL, Fagan AM, Goate A, Fox NC, et al. Clinical and biomarker changes in dominantly inherited Alzheimer’s disease. N Engl J Med. 2012;367:795–804.

[15] Nesse RM, Finch CE, Nunn CL. Does selection for short sleep duration explain human vulnerability to Alzheimer’s disease? Evol Med Public Health. 2017.

[16] Lutsey PL, Misialek JR, Mosley TH, Gottesman RF, Punjabi NM, Shahar E, et al. Sleep characteristics and risk of dementia and Alzheimer’s disease: The Atherosclerosis Risk in Communities Study. Alzheimers Dement. 2018;14:157–66.

[17] Ju YS, Ooms SJ, Sutphen C, Macauley SL, Zangrilli MA, Jerome G, et al. Slow wave sleep disruption increases cerebrospinal fluid amyloid-beta levels. Brain. 2017;140:2104–11.

[18] de Almondes KM, Costa MV, Malloy-Diniz LF, Diniz BS. Insomnia and risk of dementia in older adults: Systematic review and meta-analysis. J Psychiatr Res. 2016;77:109–15.

[19] Carvalho DZ, St Louis EK, Knopman DS, Boeve BF, Lowe VJ, Roberts RO, et al. Association of Excessive Daytime Sleepiness With Longitudinal beta-Amyloid Accumulation in Elderly Persons Without Dementia. JAMA Neurol. 2018.

[20] Faubel R, Lopez-Garcia E, Guallar-Castillon P, Graciani A, Banegas JR, Rodriguez-Artalejo F. Usual sleep duration and cognitive function in older adults in Spain. J Sleep Res. 2009;18:427–35.

[21] Sindi S, Johansson L, Skoog J, Darin Mattsson A, Sjöberg L, Wang H-X, et al. Sleep disturbances and later cognitive status: A multi-centre study. Sleep Medicine.

[22] Suh SW, Han JW, Lee JR, Byun S, Kwon SJ, Oh SH, et al. Sleep and cognitive decline: A prospective nondemented elderly cohort study. Ann Neurol. 2018;83:472–82.

[23] Diem SJ, Blackwell TL, Stone KL, Yaffe K, Tranah G, Cauley JA, et al. Measures of Sleep-Wake Patterns and Risk of Mild Cognitive Impairment or Dementia in Older Women. Am J Geriatr Psychiatry. 2016;24:248–58.

[24] Smith GD, Ebrahim S. ‘Mendelian randomization’: can genetic epidemiology contribute to understanding environmental determinants of disease? Int J Epidemiol. 2003;32:1–22.

[25] Smith GD, Lawlor DA, Harbord R, Timpson N, Day I, Ebrahim S. Clustered environments and randomized genes: a fundamental distinction between conventional and genetic epidemiology. PLoS Med. 2007;4:e352.

[26] Lawlor DA. Commentary: Two-sample Mendelian randomization: opportunities and challenges. Int J Epidemiol. 2016;45:908–15.

[27] Bowden J, Davey Smith G, Burgess S. Mendelian randomization with invalid instruments: effect estimation and bias detection through Egger regression. Int J Epidemiol. 2015;44:512–25.

[28] Jones SE, Lane JM, Wood AR, van Hees VT, Tyrrell J, Beaumont RN, et al. Genome-wide association analyses of chronotype in 697,828 individuals provides new insights into circadian rhythms in humans and links to disease. bioRxiv. 2018.

[29] Dashti H, Jones SE, Wood AR, Lane J, van Hees VT, Wang H, et al. GWAS in 446,118 European adults identifies 78 genetic loci for self-reported habitual sleep duration supported by accelerometer-derived estimates. bioRxiv. 2018.

[30] Lane JM, Jones S, Dashti H, Wood A, Aragam K, van Hees VT, et al. Biological and clinical insights from genetics of insomnia symptoms. bioRxiv. 2018.

[31] Jones SE, van Hees VT, Mazzotti DR, Marques-Vidal P, Sabia S, van der Spek A, et al. Genetic studies of accelerometer-based sleep measures in 85,670 individuals yield new insights into human sleep behaviour. bioRxiv. 2018.

[32] Doherty A, Jackson D, Hammerla N, Plotz T, Olivier P, Granat MH, et al. Large Scale Population Assessment of Physical Activity Using Wrist Worn Accelerometers: The UK Biobank Study. PLoS One. 2017;12:e0169649.

[33] Burgess S, Thompson SG, Collaboration CCG. Avoiding bias from weak instruments in Mendelian randomization studies. Int J Epidemiol. 2011;40:755–64.

[34] Staiger D, Stock JH. Instrumental Variables Regression with Weak Instruments. Econometrica. 1997;65:557–86.

[35] Lambert JC, Ibrahim-Verbaas CA, Harold D, Naj AC, Sims R, Bellenguez C, et al. Meta-analysis of 74,046 individuals identifies 11 new susceptibility loci for Alzheimer’s disease. Nat Genet. 2013;45:1452–8.

[36] Gibran Hemani JZ, Kaitlin H Wade, Charles Laurin, Benjamin Elsworth, Stephen Burgess, Jack Bowden, Ryan Langdon, Vanessa Tan, James Yarmolinsky, Hashem A. Shihab, Nicholas Timpson, David M Evans, Caroline Relton, Richard M Martin, George Davey Smith, Tom R Gaunt, Philip C Haycock. MR-Base: a platform for systematic causal inference across the phenome using billions of genetic associations. bioRxiv. 2017.

[37] Burgess S, Labrecque JA. Mendelian randomization with a binary exposure variable: interpretation and presentation of causal estimates. Eur J Epidemiol. 2018;33:947–52.

[38] Bowden J, Davey Smith G, Haycock PC, Burgess S. Consistent Estimation in Mendelian Randomization with Some Invalid Instruments Using a Weighted Median Estimator. Genet Epidemiol. 2016;40:304–14.

[39] Higgins JP, Thompson SG, Deeks JJ, Altman DG. Measuring inconsistency in meta-analyses. BMJ. 2003;327:557–60.

[40] Bowden J, Del Greco MF, Minelli C, Davey Smith G, Sheehan NA, Thompson JR. Assessing the suitability of summary data for two-sample Mendelian randomization analyses using MR-Egger regression: the role of the I2 statistic. Int J Epidemiol. 2016.

[41] Stone M. Cross-validatory choice and assessment of statistical predictions. J R Stat Soc B 1974;36:111–47.

[42] Hartwig FP, Davies NM, Hemani G, Davey Smith G. Two-sample Mendelian randomization: avoiding the downsides of a powerful, widely applicable but potentially fallible technique. Int J Epidemiol. 2016;45:1717–26.

[43] Hemani G, Tilling K, Davey Smith G. Orienting the causal relationship between imprecisely measured traits using GWAS summary data. PLoS Genet. 2017;13:e1007081.

[44] Jansen IE, Savage JE, Watanabe K, Bryois J, Williams DM, Steinberg S, et al. Genome-wide meta-analysis identifies new loci and functional pathways influencing Alzheimer’s disease risk. Nat Genet. 2019;51:404–13.

[45] Burgess S, Davies NM, Thompson SG. Bias due to participant overlap in two-sample Mendelian randomization. Genet Epidemiol. 2016;40:597–608.

[46] Zetterberg H, Alexander DM, Spandidos DA, Blennow K. Additional evidence for antagonistic pleiotropic effects of APOE. Alzheimers Dement. 2009;5:75.

[47] Hartmann JA, Carney CE, Lachowski A, Edinger JD. Exploring the construct of subjective sleep quality in patients with insomnia. J Clin Psychiatry. 2015;76:e768–73.

[48] Munafo MR, Tilling K, Taylor AE, Evans DM, Davey Smith G. Collider scope: when selection bias can substantially influence observed associations. Int J Epidemiol. 2018;47:226–35.

[49] Krokstad S, Langhammer A, Hveem K, Holmen TL, Midthjell K, Stene TR, et al. Cohort Profile: the HUNT Study, Norway. Int J Epidemiol. 2013;42:968–77.

[50] Dhand R, Sohal H. Good sleep, bad sleep! The role of daytime naps in healthy adults. Current Opinion in Pulmonary Medicine. 2006;12:379–82.

[51] Mantua J, Spencer RMC. Exploring the nap paradox: are mid-day sleep bouts a friend or foe? Sleep Med. 2017;37:88–97.

[52] Takahashi M. The role of prescribed napping in sleep medicine. Sleep Med Rev. 2003;7:227–35.

[53] Faraut B, Andrillon T, Vecchierini MF, Leger D. Napping: A public health issue. From epidemiological to laboratory studies. Sleep Med Rev. 2017;35:85–100.

[54] Chen G, Tang K, Chen F, Wei S, Lin F, Yao J, et al. Afternoon nap and nighttime sleep with risk of micro- and macrovascular disease in middle-aged and elderly population. Int J Cardiol. 2015;187:553–5.

[55] Dautovich ND, Kay DB, Perlis ML, Dzierzewski JM, Rowe MA, McCrae CS. Day-to-day variability in nap duration predicts medical morbidity in older adults. Health Psychol. 2012;31:671–6.

[56] Baran B, Mantua J, Spencer RM. Age-related Changes in the Sleep-dependent Reorganization of Declarative Memories. J Cogn Neurosci. 2016;28:792–802.

[57] Cairney SA, Guttesen AAV, El Marj N, Staresina BP. Memory Consolidation Is Linked to Spindle-Mediated Information Processing during Sleep. Curr Biol. 2018;28:948–54 e4.

[58] Ruch S, Markes O, Duss SB, Oppliger D, Reber TP, Koenig T, et al. Sleep stage II contributes to the consolidation of declarative memories. Neuropsychologia. 2012;50:2389–96.

[59] Antonenko D, Diekelmann S, Olsen C, Born J, Molle M. Napping to renew learning capacity: enhanced encoding after stimulation of sleep slow oscillations. Eur J Neurosci. 2013;37:1142–51.

[60] Brooks A, Lack L. A brief afternoon nap following nocturnal sleep restriction: which nap duration is most recuperative? Sleep. 2006;29:831–40.

[61] Cremone A, Kurdziel LBF, Fraticelli-Torres A, McDermott JM, Spencer RMC. Napping reduces emotional attention bias during early childhood. Dev Sci. 2017;20.

[62] Boyle PA, Malloy PF, Salloway S, Cahn-Weiner DA, Cohen R, Cummings JL. Executive dysfunction and apathy predict functional impairment in Alzheimer disease. Am J Geriatr Psychiatry. 2003;11:214–21.

[63] Hargrave R, Maddock RJ, Stone V. Impaired recognition of facial expressions of emotion in Alzheimer’s disease. J Neuropsychiatry Clin Neurosci. 2002;14:64–71.

[64] Fry A, Littlejohns TJ, Sudlow C, Doherty N, Adamska L, Sprosen T, et al. Comparison of Sociodemographic and Health-Related Characteristics of UK Biobank Participants With Those of the General Population. Am J Epidemiol. 2017;186:1026–34.

[65] van Hees VT, Sabia S, Jones SE, Wood AR, Anderson KN, Kivimaki M, et al. Estimating sleep parameters using an accelerometer without sleep diary. Sci Rep. 2018;8:12975.

