## Supplementary material for "Is disrupted sleep a risk factor for Alzheimer’s disease? Evidence from a two-sample Mendelian randomization analysis": Online Supplement

**Online Supplement Contents**

|  | **Supplement Item** | **Page Number** |
| --- | --- | --- |
| **Methods** | Details of sensitivity analyses for sleep traits on Alzheimer’s disease risk | 2 |
|  | Assessment of bias due to winner’s curse: independent replication sample details | 5 |
|  | Details of additional analyses for associations between genetic liability for Alzheimer’s disease and the sleep traits | 6 |
|  | Replication of associations between genetic liability for Alzheimer’s disease and frequent insomnia in the HUNT dataset | 7 |
| **Tables** | SNP flow through the harmonization procedure for the causal effects of sleep traits on Alzheimer’s disease | 8 |
|  | Phenotypic correlations of all sleep traits | 9 |
|  | Comparing causal effect estimates for sleep traits on Alzheimer’s Disease between IWV, MR-Egger and weighted median regression methods | 10 |
|  | MR-Egger intercept estimates for the causal effect of sleep traits on Alzheimer’s disease | 12 |
|  | I^2^_GX_ estimates for the causal effect of each sleep trait on Alzheimer’s disease | 13 |
|  | Between SNP heterogeneity statistics for the IVW analyses of causal effects of sleep traits on Alzheimer’s disease | 14 |
|  | Causal effect estimates for sleep traits on Alzheimer’s Disease after removing outliers detected by Radial MR | 15 |
|  | Causal effect of sleep traits on Alzheimer’s Disease after excluding palindromic SNPs | 16 |
|  | Assessment of bias due to winner’s curse: causal effect of sleep traits on Alzheimer’s Disease when using meta-analysis or replication samples | 18 |
|  | Comparing causal effect estimates for liability to Alzheimer’s Disease on sleep traits between IWV, MR-Egger and weighted median regression methods (including ApoE SNP rs429358) | 19 |
|  | Associations of ApoE SNP (rs429358) only on sleep traits | 21 |
|  | Comparing causal effect estimates for liability to Alzheimer’s Disease on sleep traits between IWV, MR-Egger and weighted median regression methods (excluding ApoE SNP rs429358) | 22 |
|  | Causal effect estimates for liability to Alzheimer’s Disease on sleep traits after removal of outliers detected by Radial MR | 24 |
|  | Associations between liability for Alzheimer’s Disease and insomnia in HUNT compared to UK Biobank (including ApoE SNP rs429358 as an instrument for Alzheimer’s disease) | 25 |
| **Figures** | Funnel plot for analysis of chronotype on AD risk | 26 |
|  | Funnel plot for analysis of daytime napping on AD risk | 27 |
|  | Funnel plot for analysis of L5 timing on AD risk | 28 |
| **References** |  | 29 |

**Sensitivity analyses**
A series of sensitivity analyses were conducted to check for violation of the key MR assumptions and check the robustness of the causal effect estimates:
(1) The IVW method assumes no horizontal pleiotropy (i.e. it assumes there are no causal paths from the SNPs to Alzheimer’s disease that do not go through the sleep trait of interest).^1^ It also assumes the gene-exposure association estimates are measured without error (the no measurement error [NOME] assumption).^1^ Thus, we compared results from the IVW regressions to those obtained with MR-Egger regressions,^1^ as the use of multiple alleles in MR analyses increases the potential for pleiotropic effects due to aggregation of invalid genetic instruments.^2^ MR-Egger assumes NOME but relaxes the assumption that the effects of genetic variants on the outcome operate entirely via the exposure (i.e. it does not assume no horizontal pleiotropy), by not constraining the intercept term to zero in the weighted regression described above. In this instance, the intercept parameter indicates the overall pleiotropic effect of the SNPs on the outcome, with a non-zero intercept providing evidence for bias due to pleiotropy. The beta coefficient (or slope) of MR-Egger provides a causal estimate of the exposure on the outcome, accounting for this level of pleiotropy and assuming that the pleiotropic effect of SNPs on the outcome is not correlated with the instrument strength (which would invalidate the previously mentioned InSIDE assumption).^1^
(2) We assessed the NOME assumption using an adaptation of the I^2^ statistic^3^, referred to as I^2^_GX._ I^2^_GX_ provides an estimate of the degree of regression dilution in the MR-Egger causal estimate due to uncertainty in the SNP-exposure estimates. We then used simulation extrapolation (SIMEX) to adjust the MR-Egger estimate for this dilution, as described previously^4^.
(3) We compared the results from IVW and MR-Egger regression to those obtained with the weighted median method,^5^ which provides a consistent estimate of causal effect if at least 50% of the genetic variants are valid instrumental variables (i.e. robustly associated with the exposure, not associated with confounding factors and only associated with the outcome via the exposure of interest).
(4) To further minimise bias due to horizontal pleiotropy, we used Radial MR methods^6^ to detect and remove any SNP outliers prior to all MR analysis being conducted. Outliers are defined as those SNPs that contribute the most heterogeneity to Cochran’s Q, based on a multiple testing corrected P value threshold.
(4) The presence of excessive between-SNP heterogeneity in an MR analysis can be an indication that some of the genetic variants are pleiotropic. Thus, we assessed heterogeneity (i.e. variability in estimates from different genetic variants) using Cochran’s Q statistic^1^. Funnel plots were then generated to enable visual assessment of the extent to which pleiotropy is likely to balanced (or directional) across the set of instruments used in each analysis. Symmetry in these plots provides evidence of no directional horizontal pleiotropy^6^.
(5) To further assess the influence of potentially pleiotropic SNPs on the causal estimates^7^ we conducted a leave-one-out permutation, in which each SNP was systematically removed from the analysis. If any SNP was having an undue influence on the overall causal effect estimate, we would expect to see some distortion in the distribution of these results.
(6) We checked that results were similar after excluding palindromic SNPs (i.e. SNPs whose alleles correspond to nucleotides that pair with each other in a DNA molecule^8^).
(7) In MR, it is assumed that the genetic instruments influence the exposure first and then the outcome, through the exposure. However, it is possible that the SNPs used to instrument sleep parameters may have a direct effect on AD risk, which then go on to effect sleep. To test that the hypothesised causal direction was correct for each SNP, we performed Steiger filtering^9^ which examines whether the SNP explains more variance in the exposure than it does in the outcome (which should be true if the hypothesised causal direction from exposure to outcome is correct).
(8) Finally, we investigated potential for bias due to ‘winner’s curse’; where the magnitude of the effect sizes for variants identified within a single discovery sample are likely to be larger than in the overall population, even if they are truly associated with the exposure. In two-sample MR, where there is little or no overlap in participants in the two samples (as is the case in this study), winner’s curse is likely to bias causal effect estimates towards the null.^2^ Assessment of winner’s curse was only possible for frequent insomnia and self-report sleep duration where the GWAS was replicated in independent samples (The Nord-Trøndelag Health Study [HUNT] and The Cohorts for Heart and Aging Research in Genomic Epidemiology [CHARGE] consortium, respectively). Details of the replication samples and winners curse analyses are in the online supplement.

**Assessing bias due to Winner’s curse using SNP-exposure estimates from independent replication samples**
*Frequent Insomnia*
The self-reported insomnia GWAS was replicated in The Nord-Trøndelag Health Study (HUNT) on a sample of 14,923 individuals experiencing frequent insomnia symptoms and 47,610 controls^10^. A total of 48 approximately independent genome wide significant loci were identified as being associated with frequent insomnia. Of the 48 SNPs, 47 were identified in the outcome dataset. No SNPs were excluded due to incompatible alleles.

*Sleep duration*
The self-reported sleep duration GWAS was replicated in The Cohorts for Heart and Aging Research in Genomic Epidemiology [CHARGE] consortium, on a sample of 47,180 from 18 studies^11^. A total of 70 SNPs approximately independent genome wide significant loci were available in CHARGE (some of which are proxies for the SNPs identified in the UK Biobank). Eight SNPs had no proxies available in the CHARGE genetic dataset. One SNP was not available in the AD outcome GWAS. Thus, a total of 69 SNPs were available for these analyses. No SNPs were excluded due to incompatible alleles.

**Additional analyses: Associations between genetic liability for AD and the sleep traits**

To examine whether the previously observed associations between AD and sleep disruption^12-16^ could be replicated in an MR framework, we tested whether genetic liability for AD was causally associated with the self-reported and accelerometer-measured sleep traits. SNP-exposure estimates were obtained for 20 independent genome-wide significant AD SNPs identified in the previously described IGAP AD GWAS meta-analysis^17^. IGAP report 21 AD SNPs, but two of them are in the ApoE region (rs429358 and rs7412). Together, tag they the ApoE haplotypes. Given these two SNPs are in linkage disequilibrium (i.e. they are not independent, R^2^ approximately 0.015), we included only the strongest of the two SNPs (rs429358) in all analyses. It is worth noting, however, that results did not change with the inclusion of the additional ApoE SNP (results available on request). The F statistic for the AD instruments was 159.61. To obtain the SNP-outcome estimates, we extracted those 20 AD SNPs from each of the sleep trait GWAS. No SNPs were excluded for being palindromic with intermediate allele frequencies. The number of SNPs included in each analysis is detailed in results tables below, and the table footnote contains information on which SNPs were removed from each analysis and for what reason. As with the main analysis of sleep traits on AD risk, SNP-exposure and SNP-outcome coefficients were combined using an inverse-variance-weighted (IVW). MR-Egger, weighted median, Radial MR and Steiger filtering were performed to assess potential violation of the MR assumptions. Analyses were conducted both with and without the ApoE variant included, as ApoE has been previously shown to be pleiotropic^18^ (which violates an MR assumption). We also examined associations of ApoE (as a single genetic instrument) with the sleep traits. Note that, as AD is a binary exposure, SNP-exposure coefficients are on the log odds scale. Thus, causal estimates for the effect of AD on sleep traits are rescaled so that they are interpreted per doubling of genetic liability for AD.

It is worth noting that there are several important limitations to these analyses. Firstly, the average age of AD diagnosis in the UK is around 75 years^19^, with the neuropsychological effects of prodromal disease detectable up to 8 years prior to a diagnosis of mild cognitive impairment (i.e. around 10-12 years prior to AD diagnosis)^20^. Thus, the majority of UK Biobank participants are likely too young (average age at recruitment 56 years^21^) to have even prodromal disease at the time the sleep traits were measured. Examining the effect of AD on sleep traits in a relatively young, healthy population (when we hypothesise that any effects of AD on sleep traits will likely be a result of increasing pathological burden) may not yield reliable results. Secondly, there is ‘healthy selection’ into the UK Biobank; only approximately 5% of those invited to participate in the study accepted the invitation^22^. This makes it unlikely that participants with an existing AD diagnosis or undiagnosed prodromal disease would have been recruited into the study.

**Replication of AD - frequent insomnia analyses in the HUNT dataset**

We replicated the association between genetic liability for AD and frequent insomnia in N=62,533 participants from The Nord-Trøndelag Health Study (HUNT)^10,23^. To do this, SNP-exposure estimates were obtained for 20 independent genome-wide significant AD SNPs identified in the previously described IGAP AD GWAS meta-analysis^17^. To obtain the SNP-outcome estimates, we identified the 20 AD SNPs in the GWAS of frequent insomnia performed in the HUNT dataset. No SNPs were excluded for being palindromic with intermediate allele frequencies.

**Table A: SNP flow through the harmonization procedure for the causal effects of sleep traits on Alzheimer’s disease**

|  | **Number of genome-wide significant loci identified in GWAS** | **N SNPs excluded as not biallelic** | **N SNPs unavailable in the outcome dataset** | | **N palindromic SNPs** | **Total number of SNPs included in main analysis** | **Total number of SNPs included in sensitivity analysis excluding palindromic SNPs** |
| --- | --- | --- | --- | --- | --- | --- | --- |
| **Self-reported measures** | | | |  |  |  |  |
| **Chronotype** | 351 | 10 | 13 | | 45 | 328 | 283 |
| **Sleep duration** | 78 | 0 | 4 | | 8 | 74 | 66 |
| **Short sleep duration** | 27 | 0 | 2 | | 3 | 25 | 22 |
| **Long sleep duration** | 8 | 0 | 1 | | 2 | 7 | 5 |
| **Frequent insomnia** | 48 | 0 | 1 | | 7 | 47 | 40 |
| **Daytime napping** | 170 | 0 | 19 | | 21 | 151 | 130 |
| **Daytime sleepiness** | 37 | 0 | 1 | | 3 | 36 | 33 |
| **Accelerometer measures** | | | | | | | |
| **L5 timing** | 6 | 0 | 0 | | 1 | 6 | 5 |
| **Sleep duration** | 10 | 0 | 1 | | 2 | 9 | 7 |
| **Sleep fragmentation** | 21 | 0 | 1 | | 0 | 19* | - |

Note that one of the sleep fragmentation SNPs was rs429358 in the ApoE region. As ApoE is the strongest genetic instrument for Alzheimer’s disease (the outcome in these analyses), this SNP was excluded as an instrument for sleep fragmentation due to potential for horizontal pleiotropy.

**Table B: Phenotypic correlations of all sleep traits (sample sizes for correlations are shown in parentheses)**

|  |  | **Self-reported measures** | | | | | | | | **Accelerometer measures** | | |
| --- | --- | --- | --- | --- | --- | --- | --- | --- | --- | --- | --- | --- |
|  |  | **Chronotype** | **Frequent insomnia** | **Daytime napping** | **Daytime sleepiness** | **Sleep duration** | **Short sleep duration** | **Long sleep duration** | **L5 timing** | | **Sleep duration** | **Sleep fragmentation** |
| **Self-reported measures** | **Chronotype** | **1.00 (378,597)** |  |  |  |  |  |  |  | |  |  |
|  | **Frequent Insomnia** | 0.004 (198,208) | **1.00 (198,796)** |  |  |  |  |  |  | |  |  |
|  | **Daytime napping** | 0.02 (378,446) | 0.07 (198,725) | **1.00 (379,580)** |  |  |  |  |  | |  |  |
|  | **Daytime sleepiness** | -0.004 (377,323) | 0.09 (198,113) | 0.16 (378,301) | **1.00 (378,416)** |  |  |  |  | |  |  |
|  | **Sleep duration** | -0.03 (375,591) | -0.30 (197,406) | 0.08 (377,547) | 0.002 (376,488) | **1.00 (377,695)** |  |  |  | |  |  |
|  | **Short sleep duration** | 0.01 (347,690) | 0.32 (182,747) | -0.005 (348,582) | 0.06 (347,661) | -0.82 (348,717) | **1.00 (348,717)** |  |  | |  |  |
|  | **Long sleep duration** | -0.03 286, 670) | -0.01 (138,479) | 0.10 (287,361) | 0.08  (286, 650) | 0.74 (287,456) | 0.00 (258,612) | **1.00 (287,456)** |  | |  |  |
| **Accelerometer measures** | **L5 timing** | 0.29 (77,038) | 0.009 (40,428) | 0.01 (77,278) | 0.01 (77,189) | 0.009 (77,115) | 0.005 (72,269) | 0.02 (60,614) | **1.00 (77,287)** | |  |  |
|  | **Sleep duration** | 0.02 (77,512) | 0.04 (40,665) | -0.02 (77,757) | -0.03 (77,666) | 0.15 (77,590) | -0.10 (72,721) | 0.09 (60,968) | -0.001 (76,728) | | **1.00 (77,766)** |  |
|  | **Sleep fragmentation** | -0.005 (77,512) | 0.01 (40,665) | 0.02 (77,757) | -0.01 (77,666) | 0.08 (77,590) | -.0.04 (72,721) | 0.05 (60,968) | -0.007 (76,728) | | 0.28 (77,766) | **1.00 (77,766)** |

**Table C: Comparing causal effect estimates for sleep traits on Alzheimer’s Disease between IWV, MR-Egger and weighted median regression methods**

|  |  | **Self-reported measures** | | |  | **Accelerometer measures** | | |
| --- | --- | --- | --- | --- | --- | --- | --- | --- |
| **Sleep trait** | **Units of exposure** | **N SNPs** | **OR (95% CI)** | **P** |  | **N SNPs** | **OR (95% CI)** | **P** |
| **Chronotype** | Per category increase from (i) definitely a morning person, (ii) more a morning than an evening person, (iii) do not know, (iv) more an evening than morning person and (vi) definitely an evening person |  |  |  |  |  |  |  |
| IVW |  | 328 | 0.98 (0.86, 1.11) | 0.77 |  |  |  |  |
| MR-Egger |  |  | 0.94 (0.63, 1.41) | 0.76 |  |  |  |  |
| SIMEX corrected MR-Egger |  |  | 0.92 (0.56, 1.52) | 0.73 |  |  |  |  |
| Weighted Median |  |  | 1.02 (0.85, 1.22) | 0.84 |  |  |  |  |
| **L5 timing** | Per hour elapsed since previous midnight |  |  |  |  |  |  |  |
| IVW |  |  |  |  |  | 6 | 0.73 (0.40, 1.31) | 0.29 |
| MR-Egger |  |  |  |  |  |  | 0.34 (0.12, 0.96) | 0.11 |
| SIMEX corrected MR-Egger |  |  |  |  |  |  | 0.32 (0.11, 0.93) | 0.10 |
| Weighted Median |  |  |  |  |  |  | 0.94 (0.55, 1.60) | 0.82 |
| **Sleep duration** | Per hour increase |  |  |  |  |  |  |  |
| IVW |  | 74 | 0.94 (0.71, 1.25) | 0.66 |  | 9 | 0.74 (0.48, 1.16) | 0.19 |
| MR-Egger |  |  | 0.46 (0.15, 1.39) | 0.17 |  |  | 2.32 (0.83, 6.51) | 0.15 |
| SIMEX corrected MR-Egger |  |  | 0.34 (0.08, 1.46) | 0.15 |  |  | 2.48 (0.85, 7.25) | 1.14 |
| Weighted Median |  |  | 0.82 (0.54, 1.23) | 0.33 |  |  | 0.72 (0.43, 1.2) | 0.21 |
| **Short sleep duration** | Per doubling of genetic liability for short sleep duration |  |  |  |  |  |  |  |
| IVW |  | 25 | 1.19 (0.93, 1.53) | 0.16 |  |  |  |  |
| MR-Egger |  |  | 1.02 (0.38, 2.79) | 0.96 |  |  |  |  |
| SIMEX corrected MR-Egger |  |  | 1.05 (0.26, 4.21) | 0.94 |  |  |  |  |
| Weighted Median |  |  | 1.20 (0.88, 1.63) | 0.25 |  |  |  |  |
| **Long sleep duration** |  |  |  |  |  |  |  |  |
| IVW |  |  | 1.10 (0.79, 1.54) | 0.57 |  |  |  |  |
| MR-Egger | Per doubling of genetic liability for long sleep duration | 7 | 1.52 (0.44, 5.20) | 0.54 |  |  |  |  |
| SIMEX corrected MR-Egger |  |  | 1.75 (0.38, 8.11) | 0.51 |  |  |  |  |
| Weighted Median |  |  | 0.87 (0.61, 1.25) | 0.45 |  |  |  |  |
| **Frequent insomnia** | Per doubling of genetic liability for frequent insomnia |  |  |  |  |  |  |  |
| IVW |  |  | 1.00 (0.59, 1.68) | 0.99 |  |  |  |  |
| MR-Egger |  | 47 | 0.57 (0.07, 4.94) | 0.61 |  |  |  |  |
| SIMEX corrected MR-Egger |  |  | 0.42 (0.03, 5.78) | 0.52 |  |  |  |  |
| Weighted Median |  |  | 1.87 (0.87, 4.04) | 0.11 |  |  |  |  |
| **Daytime napping** |  |  |  |  |  |  |  |  |
| IVW | Per category increase from (i) never, (ii) sometimes and (iii) usually | 151 | 0.70 (0.50, 0.99) | 0.04 |  |  |  |  |
| MR-Egger |  |  | 0.55 (0.16, 1.89) | 0.34 |  |  |  |  |
| SIMEX corrected MR-Egger |  |  | 0.26 (0.05, 1.22) | 0.09 |  |  |  |  |
| Weighted Median |  |  | 0.67 (0.42, 1.07) | 0.10 |  |  |  |  |
| **Daytime sleepiness** | Per category increase from (i) never or rare), (ii) sometimes, (iii) often and (iv) all the time |  |  |  |  |  |  |  |
| IVW |  | 36 | 0.65 (0.29, 1.43) | 0.28 |  |  |  |  |
| MR-Egger |  |  | 0.09 (0.002, 3.45) | 0.20 |  |  |  |  |
| SIMEX corrected MR-Egger |  |  | 0.03 (0.0002, 4.31) | 0.18 |  |  |  |  |
| Weighted Median |  |  | 0.67 (0.21, 2.13) | 0.50 |  |  |  |  |
| **Sleep fragmentation** | Per one unit increase in the number of sleep episodes |  |  |  |  |  |  |  |
| IVW |  |  |  |  |  | 19 | 1.12 (0.87, 1.45) | 0.37 |
| MR-Egger |  |  |  |  |  |  | 0.52 (0.17, 1.59) | 0.27 |
| SIMEX corrected MR-Egger |  |  |  |  |  |  | 0.41 (0.09, 1.75) | 0.24 |
| Weighted Median |  |  |  |  |  |  | 1.11 (0.78, 1.57) | 0.57 |

Grey shaded areas mean that there is no data available to conduct these analyses

**Table D. MR-Egger intercept estimates for the causal effect of sleep traits on Alzheimer’s disease**

|  | **Self-reported measures** | | | |  | **Accelerometer measures** | | | |
| --- | --- | --- | --- | --- | --- | --- | --- | --- | --- |
|  | **N SNPs** | **Log(OR)** | **SE** | **P** |  | **N SNPs** | **Log(OR)** | **SE** | **P** |
| **Chronotype** | 167 | 0.001 | 0.006 | 0.85 |  |  |  |  |  |
| **L5 timing** |  |  |  |  |  | 5 | 0.02 | 0.02 | 0.31 |
| **Sleep Duration** | 74 | 0.01 | 0.01 | 0.20 |  | 9 | -0.04 | 0.02 | 0.05 |
| **Short sleep duration** | 25 | 0.005 | 0.02 | 0.76 |  |  |  |  |  |
| **Long sleep duration** | 7 | -0.02 | 0.04 | 0.62 |  |  |  |  |  |
| **Frequent insomnia** | 47 | 0.006 | 0.01 | 0.60 |  |  |  |  |  |
| **Daytime napping** | 151 | 0.002 | 0.006 | 0.68 |  |  |  |  |  |
| **Daytime sleepiness** | 36 | 0.01 | 0.01 | 0.28 |  |  |  |  |  |
| **Sleep fragmentation** |  |  |  |  |  | 17 | 0.02 | 0.02 | 0.21 |

Grey shaded areas indicate where no data is available to conduct these analyses. OR- Odds ratio. SE – Standard error. P – P value.

**Table E. I^2^_GX_ estimates for the causal effect of each sleep trait on Alzheimer’s disease**

|  | **Self-report measures** |  | **Accelerometer measures** |
| --- | --- | --- | --- |
|  | **I^2^_GX_** |  | **I^2^_GX_** |
| **Chronotype** | 0.61 |  |  |
| **L5 timing** |  |  | 0.70 |
| **Sleep Duration** | 0.43 |  | 0.38 |
| **Short sleep duration** | 0 |  |  |
| **Long sleep duration** | 0.54 |  |  |
| **Frequent insomnia** | 0.21 |  |  |
| **Daytime napping** | 0.72 |  |  |
| **Daytime sleepiness** | 0.23 |  |  |
| **Sleep fragmentation** |  |  | 0 |

Grey shaded areas indicate where no data is available to conduct these analyses

**Table F. Between SNP heterogeneity statistics for the IVW analyses of causal effects of sleep traits on Alzheimer’s disease**

|  | **Self-reported measures** | | | |  | **Accelerometer measures** | | | |
| --- | --- | --- | --- | --- | --- | --- | --- | --- | --- |
|  | **N SNPs** | **Q** | **df** | **P** |  | **N SNPs** | **Q** | **df** | **P** |
| **Chronotype** | 328 | 218.6 | 327 | 0.001 |  |  |  |  |  |
| **L5 timing** |  |  |  |  |  | 6 | 11.7 | 5 | 0.04 |
| **Sleep Duration** | 74 | 89.65 | 73 | 0.09 |  | 9 | 11.38 | 8 | 0.18 |
| **Short sleep duration** | 25 | 33.52 | 24 | 0.09 |  |  |  |  |  |
| **Long sleep duration** | 7 | 11.62 | 6 | 0.07 |  |  |  |  |  |
| **Frequent insomnia** | 47 | 35.45 | 46 | 0.81 |  |  |  |  |  |
| **Daytime napping** | 151 | 183 | 150 | 0.03 |  |  |  |  |  |
| **Daytime sleepiness** | 36 | 35.99 | 35 | 0.42 |  |  |  |  |  |
| **Sleep fragmentation** |  |  |  |  |  | 17 | 12.01 | 16 | 0.74 |

Grey shaded areas indicate where no data is available to conduct these analyses

**Table G: Causal effect estimates for sleep traits on Alzheimer’s Disease after removing outliers detected by Radial MR.**

|  |  | **Accelerometer measures** | | |
| --- | --- | --- | --- | --- |
| **Sleep trait** | **Units of exposure** | **N SNPs** | **OR (95% CI)** | **P** |
| **L5 timing** | Per hour elapsed since previous midnight |  |  |  |
| IVW |  | 5 | 0.92 (0.61, 1.39) | 0.69 |
| MR-Egger |  |  | 0.57 (0.23, 1.38) | 0.30 |
| SIMEX corrected MR-Egger |  |  | 0.32 (0.11, 0.95) | 0.11 |
| Weighted Median |  |  | 0.98 (0.60, 1.60) | 0.94 |
| **Sleep fragmentation** | Per one unit increase in the number of sleep episodes |  |  |  |
| IVW |  | 17 | 1.00 (0.76, 1.31) | 0.99 |
| MR-Egger |  |  | 0.47 (0.15, 1.51) | 0.23 |
| SIMEX corrected MR-Egger |  |  | 0.35 (0.01, 1.26) | 0.13 |
| Weighted Median |  |  | 1.09 (0.76, 1.56) | 0.65 |

**Table H: Causal effect of sleep traits on Alzheimer’s Disease after excluding palindromic SNPs**

|  |  | **Self-reported measures** | | |  | **Accelerometer measures** | | |
| --- | --- | --- | --- | --- | --- | --- | --- | --- |
| **Sleep trait** | **Units of exposure** | **N SNPs** | **B (95% CI)** | **P** |  | **N SNPs** | **B (95% CI)** | **P** |
| **Chronotype** | Per category increase from (i) definitely a morning person, (ii) more a morning than an evening person, (iii) do not know, (iv) more an evening than morning person and (vi) definitely an evening person |  |  |  |  |  |  |  |
| IVW |  | 283 | 0.98 (0.85, 1.12) | 0.74 |  |  |  |  |
| MR-Egger |  |  | 0.89 (0.58, 1.37) | 0.61 |  |  |  |  |
| Weighted Median |  |  | 1.00 (0.82, 1.20) | 0.97 |  |  |  |  |
| **L5 timing** | Per hour elapsed since previous midnight |  |  |  |  |  |  |  |
| IVW |  |  |  |  |  | 5 | 0.92 (0.61, 1.39) | 0.69 |
| MR-Egger |  |  |  |  |  |  | 0.57 (0.23, 1.38) | 0.30 |
| Weighted Median |  |  |  |  |  |  | 0.98 (0.61, 1.59) | 0.94 |
| **Sleep duration** | Per hour increase |  |  |  |  |  |  |  |
| IVW |  | 66 | 1.00 (0.99, 1.00) | 0.63 |  | 7 | 0.80 (0.48, 1.34) | 0.40 |
| MR-Egger |  |  | 0.99 (0.97, 1.01) | 0.32 |  |  | 2.12 (0.67, 6.75) | 0.26 |
| Weighted Median |  |  | 1.00 (0.99, 1.00) | 0.29 |  |  | 0.72 (0.41, 1.26) | 0.25 |
| **Short sleep duration** | Per doubling of genetic liability for short sleep duration |  |  |  |  |  |  |  |
| IVW |  | 22 | 1.15 (0.9, 1.47) | 0.26 |  |  |  |  |
| MR-Egger |  |  | 1.61 (0.57, 4.56) | 0.38 |  |  |  |  |
| Weighted Median |  |  | 1.19 (0.86, 1.65) | 0.30 |  |  |  |  |
| **Long sleep duration** | Per doubling of genetic liability for long sleep duration |  |  |  |  |  |  |  |
| IVW |  |  | 0.97 (0.67, 1.39) | 0.85 |  |  |  |  |
| MR-Egger |  | 5 | 1.17 (0.33, 4.19) | 0.82 |  |  |  |  |
| Weighted Median |  |  | 0.85 (0.58, 1.24) | 0.40 |  |  |  |  |
| **Frequent insomnia** | Per doubling of genetic liability for frequent insomnia |  |  |  |  |  |  |  |
| IVW |  |  | 1.00 (0.57, 1.78) | 0.99 |  |  |  |  |
| MR-Egger |  | 40 | 0.46 (0.05, 4.28) | 0.50 |  |  |  |  |
| Weighted Median |  |  | 1.73 (0.75, 3.97) | 0.20 |  |  |  |  |
| **Daytime napping** | Per category increase from (i) never, (ii) sometimes and (iii) usually |  |  |  |  |  |  |  |
| IVW |  | 130 | 0.71 (0.5, 1.02) | 0.06 |  |  |  |  |
| MR-Egger |  |  | 0.56 (0.16, 1.99) | 0.37 |  |  |  |  |
| Weighted Median |  |  | 0.63 (0.39, 1.02) | 0.06 |  |  |  |  |
| **Daytime sleepiness** | Per category increase from (i) never or rare), (ii) sometimes, (iii) often and (iv) all the time |  |  |  |  |  |  |  |
| IVW |  | 33 | 0.65 (0.28, 1.50) | 0.41 |  |  |  |  |
| MR-Egger |  |  | 0.04 (0.01, 2.01) | 0.70 |  |  |  |  |
| Weighted Median |  |  | 0.68 (0.21, 2.16) | 0.77 |  |  |  |  |
| **Sleep fragmentation** | Per one unit increase in the number of sleep episodes |  |  |  |  | No palindromic SNPs so results are the same as main analysis | | |
| IVW |  |  |  |  |  |  |  |  |
| MR-Egger |  |  |  |  |  |  |  |  |
| Weighted Median |  |  |  |  |  |  |  |  |

Grey shaded areas mean that there is no data available to conduct these analyses

**Table I: Assessment of bias due to winner’s curse: causal effect of sleep traits on Alzheimer’s Disease when using meta-analysis or replication samples**

|  | **Self-report measures** | | |
| --- | --- | --- | --- |
|  | **N SNPs** | **B (95% CI)** | **P** |
| **Sleep Duration (Average hours per night)** | 69* | 1.00 (1.00, 1.01) | 0.72 |
| **Frequent Insomnia (Yes/No)** | 47 | 0.99 (0.82,1.18) | 0.87 |

*8 of the SNPs identified in the UK Biobank sleep duration GWAS were not available in the CHARGE dataset and no suitable proxies could be found. One of the proxies identified in charge was not available in the outcome dataset.

**Table J: Comparing causal effect estimates for liability to Alzheimer’s Disease on sleep traits between IWV, MR-Egger and weighted median regression methods (including ApoE SNP rs429358)**

|  | **Self-reported measures** | | |  | **Accelerometer measures** | | |
| --- | --- | --- | --- | --- | --- | --- | --- |
|  | **N SNPs** | **OR (95% CI)** | **P** |  | **N SNPs** | **OR (95% CI)** | **P** |
| **Short sleep duration (<6 hours)** |  |  |  |  |  |  |  |
| IVW | 20 | 1.001 (0.999, 1.003) | 0.25 |  |  |  |  |
| MR-Egger |  | 1.001 (0.999, 1.003) | 0.33 |  |  |  |  |
| Weighted Median |  | 1.001 (0.999, 1.003) | 0.28 |  |  |  |  |
| **Long sleep duration (>9 hours)** |  |  |  |  |  |  |  |
| IVW |  | 0.998 (0.996, 0.999) | 5.44E-03 |  |  |  |  |
| MR-Egger | 20 | 0.999 (0.996, 1.0001) | 7.38E-02 |  |  |  |  |
| Weighted Median |  | 0.999 (0.996, 0.999) | 6.49E-03 |  |  |  |  |
| **Frequent insomnia (Yes/Never or rarely)** |  |  |  |  |  |  |  |
| IVW |  | 0.996 (0.991, 0.996) | 4.40E-06 |  |  |  |  |
| MR-Egger | 20 | 0.996 (0.991, 0.997) | 2.23E-03 |  |  |  |  |
| Weighted Median |  | 0.996 (0.991, 0.996) | 3.20E-06 |  |  |  |  |
| **Daytime napping (Yes/No)** |  |  |  |  |  |  |  |
| IVW | 20 | 0.996 (0.992, 0.998) | 4.12E-04 |  |  |  |  |
| MR-Egger |  | 0.996 (0.991, 0.998) | 9.51E-03 |  |  |  |  |
| Weighted Median |  | 0.996 (0.992, 0.997) | 4.82E-05 |  |  |  |  |
| **Daytime sleepiness (Yes/No)** |  |  |  |  |  |  |  |
| IVW | 20 | 0.999 (0.997, 1.001) | 0.37 |  |  |  |  |
| MR-Egger |  | 0.999 (0.996, 1.001) | 0.24 |  |  |  |  |
| Weighted Median |  | 0.999 (0.997, 1.001) | 0.27 |  |  |  |  |
|  | **N SNPs** | **B (95% CI)** | **P** |  | **N SNPs** | **B (95% CI)** | **P** |
| **Chronotype (More morningness to more eveningness)** |  |  |  |  |  |  |  |
| IVW | 20 | -0.01 (-0.00002, -0.02) | 4.96E-02 |  |  |  |  |
| MR-Egger |  | -0.01 (-0.004, -0.02) | 1.67E-02 |  |  |  |  |
| Weighted Median |  | -0.01 (-0.01, -0.02) | 7.30E-06 |  |  |  |  |
| **L5 timing (timing of least active 5 hours of the day)** |  |  |  |  |  |  |  |
| IVW |  |  |  |  | 20 | -0.01 (-0.03, -0.01) | 6.55E-05 |
| MR-Egger |  |  |  |  |  | -0.02 (-0.04, -0.01) | 6.27E-04 |
| Weighted Median |  |  |  |  |  | -0.02 (-0.03, -0.01) | 6.60E-06 |
| **Sleep duration (Average hours per night)** |  |  |  |  |  |  |  |
| IVW | 20 | -0.005 (-0.01, -0.001) | 9.86E-03 |  | 19* | -0.01 (-0.03, -0.01) | 3.19E-04 |
| MR-Egger |  | -0.005 (-0.01, 0.00003) | 6.72E-02 |  |  | -0.02 (-0.04, -0.01) | 3.27E-03 |
| Weighted Median |  | -0.005 (-0.01, -0.002) | 1.98E-03 |  |  | -0.02 (-0.03, -0.01) | 2.33E-04 |
| **Sleep fragmentation (Number of sleep periods)** |  |  |  |  |  |  |  |
| IVW |  |  |  |  | 20 | -0.02 (-0.03, -0.01) | 2.47E-04 |
| MR-Egger |  |  |  |  |  | -0.02 (-0.04, -0.01) | 7.96E-04 |
| Weighted Median |  |  |  |  |  | -0.02 (-0.04, -0.02) | 1.00E-07 |

All effect estimates are interpreted per doubling of genetic liability for AD. Grey shaded areas mean that there is no data available to conduct these analyses. *One SNP (rs6656401) was removed due to incompatible alleles

**Table K. Associations of ApoE SNP (rs429358) only on sleep traits (N SNPs = 1)**

|  | **Self-report** | |  | **Accelerometer-measured** | |
| --- | --- | --- | --- | --- | --- |
|  | **OR (95% CI)** | **P** |  | **OR (95% CI)** | **P** |
| **Short sleep duration** | 1.001 (0.999, 1.003) | 0.29 |  |  |  |
| **Long sleep duration** | 0.999 (0.996, 0.999) | 5.13E-01 |  |  |  |
| **Frequent insomnia (Yes/No)** | 0.996 (0.991, 0.996) | 7.38E-06 |  |  |  |
| **Daytime sleepiness (Yes/No)** | 0.999 (0.997, 1.001) | 0.28 |  |  |  |
| **Daytime napping** | 0.996 (0.992, 0.997) | 2.63E-05 |  |  |  |
|  | **B (95% CI)** | **P** |  | **B (95% CI)** | **P** |
| **Chronotype (Morningness vs eveningness)** | -0.01 (-0.01, -0.02) | 6.69E-06 |  |  |  |
| **L5 timing** |  |  |  | -0.02 (-0.03, -0.01) | 5.38E-06 |
| **Sleep duration (Average hours per night)** | -0.01 (-0.01, -0.003) | 9.44E-02 |  | -0.02 (-0.03, -0.01) | 5.38E-02 |
| **Sleep fragmentation (Average number of sleep periods)** |  |  |  | -0.03 (-0.04, -0.02) | 4.47E-08 |

All effect estimates are interpreted per doubling of genetic liability for AD. Grey shaded areas mean that there is no data available to conduct these analyses.

**Table L: Comparing causal effect estimates for liability to Alzheimer’s Disease on sleep traits between IWV, MR-Egger and weighted median regression methods (excluding ApoE SNP rs429358)**

|  | **Self-reported measures** | | |  | **Accelerometer measures** | | |
| --- | --- | --- | --- | --- | --- | --- | --- |
|  | **N SNPs** | **OR (95% CI)** | **P** |  | **N SNPs** | **OR (95% CI)** | **P** |
| **Short sleep duration (<6 hours)** |  |  |  |  |  |  |  |
| IVW | 19 | 1.002 (0.998, 1.004) | 0.65 |  |  |  |  |
| MR-Egger |  | 1.007 (0.993, 1.012) | 0.76 |  |  |  |  |
| Weighted Median |  | 1.003 (0.997, 1.005) | 0.77 |  |  |  |  |
| **Long sleep duration (>9 hours)** |  |  |  |  |  |  |  |
| IVW |  | 1.002 (0.994, 0.999) | 9.18E-02 |  |  |  |  |
| MR-Egger | 19 | 1.007 (0.990, 1.008) | 0.83 |  |  |  |  |
| Weighted Median |  | 1.03 (0.995, 1.002) | 0.67 |  |  |  |  |
| **Frequent insomnia (Yes/Never or rarely)** |  |  |  |  |  |  |  |
| IVW |  | 1.003 (0.989, 0.998) | 5.50E-02 |  |  |  |  |
| MR-Egger | 19 | 1.01 (0.984, 1.01) | 0.96 |  |  |  |  |
| Weighted Median |  | 1.004 (0.986, 0.999) | 0.12 |  |  |  |  |
| **Daytime napping (Yes/No)** |  |  |  |  |  |  |  |
| IVW | 19 | 1.004 (0.991, 1.0004) | 0.21 |  |  |  |  |
| MR-Egger |  | 1.012 (0.983, 1.015) | 0.91 |  |  |  |  |
| Weighted Median |  | 1.004 (0.992, 1.003) | 0.50 |  |  |  |  |
| **Daytime sleepiness (Yes/No)** |  |  |  |  |  |  |  |
| IVW | 19 | 1.002 (0.176, 5.679) | 0.89 |  |  |  |  |
| MR-Egger |  | 1.008 (0.398, 2.485) | 0.47 |  |  |  |  |
| Weighted Median |  | 1.003 (0.148, 6.775) | 0.98 |  |  |  |  |
| **Chronotype (More morningness to more eveningness)** |  |  |  |  |  |  |  |
| IVW | 19 | 0.004 (-0.01, 0.03) | 0.55 |  |  |  |  |
| MR-Egger |  | -0.008 (-0.08, 0.06) | 0.76 |  |  |  |  |
| Weighted Median |  | -0.005 (-0.03, 0.01) | 0.49 |  |  |  |  |
| **L5 timing (timing of least active 5 hours of the day)** |  |  |  |  |  |  |  |
| IVW |  |  |  |  | 19 | -0.01 (-0.03, 0.02) | 0.55 |
| MR-Egger |  |  |  |  |  | -0.03 (-0.10, 0.05) | 0.45 |
| Weighted Median |  |  |  |  |  | -0.02 (-0.05, 0.02) | 0.35 |
| **Sleep duration (Average hours per night)** |  |  |  |  |  |  |  |
| IVW | 19 | -0.006 (-0.02, 0.006) | 0.25 |  | 18* | -0.01 (-0.04, 0.01) | 0.18 |
| MR-Egger |  | 0.0006 (-0.05, 0.05) | 0.97 |  |  | -0.06 (-0.15, -0.01) | 3.01E-02 |
| Weighted Median |  | 0.00 (-0.01, 0.02) | 0.78 |  |  | -0.01 (-0.05, 0.01) | 0.19 |
| **Sleep fragmentation (Number of sleep periods)** |  |  |  |  |  |  |  |
| IVW |  |  |  |  | 19 | 0.01 (-0.02, 0.02) | 0.89 |
| MR-Egger |  |  |  |  |  | 0.05 (-0.07, 0.05) | 0.80 |
| Weighted Median |  |  |  |  |  | 0.02 (-0.03, 0.009) | 0.43 |

All effect estimates are interpreted per doubling of genetic liability for AD. Grey shaded areas mean that there is no data available to conduct these analyses. *One SNP (rs6656401) was removed due to incompatible alleles

**Table M: Causal effect estimates for liability to Alzheimer’s Disease on sleep traits after removal of outliers detected by Radial MR**

|  | **Self-reported measures** | | |  | **Accelerometer measures** | | |
| --- | --- | --- | --- | --- | --- | --- | --- |
|  | **N SNPs** | **B (95% CI)** | **P** |  | **N SNPs** | **B (95% CI)** | **P** |
| **Chronotype (More morningness to more eveningness)** |  |  |  |  |  |  |  |
| IVW | 17**** | -0.01 (-0.004, -0.02) | 2.67E-01 |  |  |  |  |
| MR-Egger |  | -0.01 (-0.01, -0.02) | 1.72E-02 |  |  |  |  |
| Weighted Median |  | -0.01 (-0.01, -0.02) | 3.60E-06 |  |  |  |  |
| **Sleep duration (Average hours per night)** |  |  |  |  |  |  |  |
| IVW | 19** | -0.01 (-0.01, -0.002) | 4.40E-03 |  | 18* | -0.01 (-0.03, -0.009) | 7.77E-05 |
| MR-Egger |  | -0.01 (-0.01, -0.002) | 2.19E-02 |  |  | -0.01 (-0.03, -0.007) | 4.21E-03 |
| Weighted Median |  | -0.005 (-0.01, -0.003) | 1.65E-03 |  |  | -0.01 (-0.03, -0.007) | 4.95E-04 |
| **Sleep fragmentation (Number of sleep periods)** |  |  |  |  |  |  |  |
| IVW |  |  |  |  | 19*** | -0.02 (-0.03, -0.01) | 4.10E-06 |
| MR-Egger |  |  |  |  |  | -0.02 (-0.04, -0.01) | 2.99E-06 |
| Weighted Median |  |  |  |  |  | -0.02 (-0.03, -0.02) | 1.00E-07 |

All effect estimates are interpreted per doubling of genetic liability for AD. Grey shaded areas mean that there is no data available to conduct these analyses.
*One SNP (rs6656401) was removed due to incompatible alleles and another SNP (rs1476679) was removed due to being detected as an outlier with Radial MR.
** rs10838725 was removed due to being detected as an outlier with Radial MR.
*** rs9271192 was removed due to being detected as an outlier with Radial MR.

**** rs4147929, rs190982 and rs10838725 were removed due to being detected as outliers with Radial MR.

**Table N: Associations between liability for Alzheimer’s Disease and insomnia in HUNT compared to UK Biobank (including ApoE SNP rs429358 as an instrument for Alzheimer’s disease)**

|  | **N for outcome sample** | **N SNPs** | **OR (95% CI)** | **P** |
| --- | --- | --- | --- | --- |
| **HUNT insomnia** |  |  |  |  |
| IVW |  | 20 | 0.979 (0.965, 1.006) | 0.17 |
| MR-Egger | 62, 533 |  | 0.970 (0.954, 1.004) | 0.11 |
| Weighted Median |  |  | 0.977 (0.967, 1.002) | 0.09 |
| **UK Biobank insomnia** |  |  |  |  |
| IVW |  |  | 0.998 (0.996, 0.999) | 5.44E-03 |
| MR-Egger | 453,379 | 20 | 0.999 (0.996, 1.0001) | 7.38E-02 |
| Weighted Median |  |  | 0.999 (0.996, 0.999) | 6.49E-03 |

All effect estimates are interpreted per doubling of genetic liability for AD.

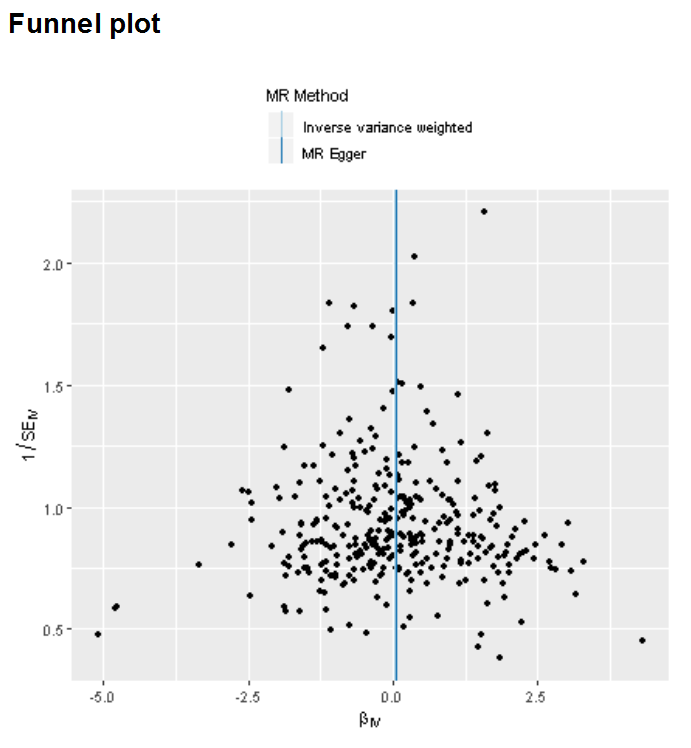

**Figure A – Funnel plot for analysis of chronotype on AD risk**

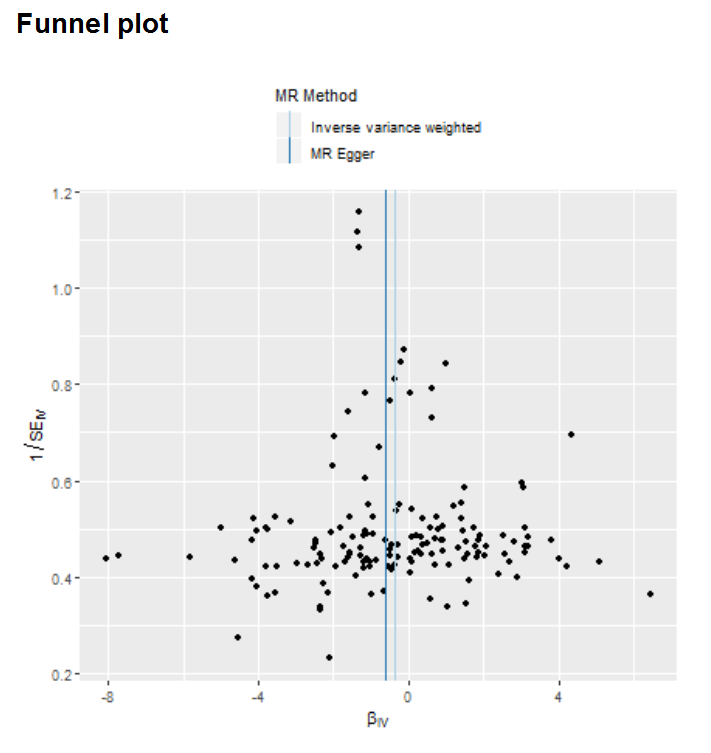

**Figure B – Funnel plot for analysis of daytime napping on AD risk**

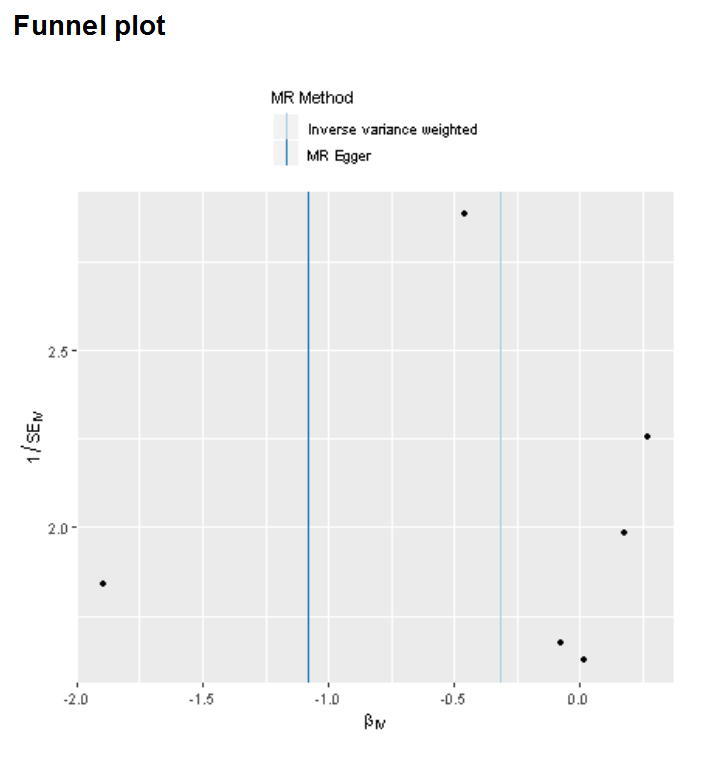

**Figure C – Funnel plot for analysis of L5 timing on AD risk**

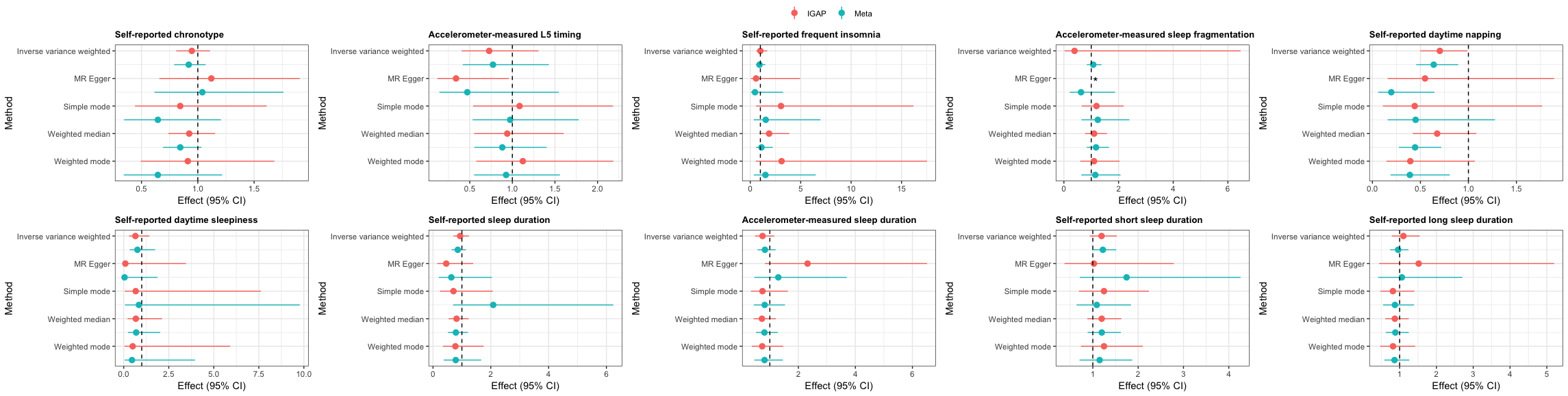

**Figure D – Comparing results for associations of sleep traits on risk of AD when using IGAP GWAS alone (n=17,008 AD cases and 37,154 controls; IGAP) vs the GWAS meta-analysis of IGAP, the AD working group of the Psychiatric Genomics Consortium and the AD Sequencing Project (n= 24,087 cases and 55,058 controls; Meta)**

***MR Egger estimate for effect of accelerometer-measured sleep fragmentation not plotted due to imprecision**

**References**

1.	Bowden J, Davey Smith G, Burgess S. Mendelian randomization with invalid instruments: effect estimation and bias detection through Egger regression. *Int J Epidemiol.* 2015;44(2):512-525.

2.	Haycock PC, Burgess S, Wade KH, Bowden J, Relton C, Davey Smith G. Best (but oft-forgotten) practices: the design, analysis, and interpretation of Mendelian randomization studies. *Am J Clin Nutr.* 2016;103(4):965-978.

3.	Higgins JP, Thompson SG, Deeks JJ, Altman DG. Measuring inconsistency in meta-analyses. *BMJ.* 2003;327(7414):557-560.

4.	Bowden J, Del Greco MF, Minelli C, Davey Smith G, Sheehan NA, Thompson JR. Assessing the suitability of summary data for two-sample Mendelian randomization analyses using MR-Egger regression: the role of the I2 statistic. *Int J Epidemiol.* 2016.

5.	Bowden J, Davey Smith G, Haycock PC, Burgess S. Consistent Estimation in Mendelian Randomization with Some Invalid Instruments Using a Weighted Median Estimator. *Genet Epidemiol.* 2016;40(4):304-314.

6.	Bowden J, Spiller W, Del Greco MF, et al. Improving the visualization, interpretation and analysis of two-sample summary data Mendelian randomization via the Radial plot and Radial regression. *Int J Epidemiol.* 2018.

7.	Stone M. Cross-validatory choice and assessment of statistical predictions. *J R Stat Soc B* 1974;36:111-147.

8.	Hartwig FP, Davies NM, Hemani G, Davey Smith G. Two-sample Mendelian randomization: avoiding the downsides of a powerful, widely applicable but potentially fallible technique. *Int J Epidemiol.* 2016;45(6):1717-1726.

9.	Hemani G, Tilling K, Davey Smith G. Orienting the causal relationship between imprecisely measured traits using GWAS summary data. *PLoS Genet.* 2017;13(11):e1007081.

10.	Lane JM, Jones S, Dashti H, et al. Biological and clinical insights from genetics of insomnia symptoms. *bioRxiv.* 2018.

11.	Gottlieb DJ, Hek K, Chen TH, et al. Novel loci associated with usual sleep duration: the CHARGE Consortium Genome-Wide Association Study. *Mol Psychiatry.* 2015;20(10):1232-1239.

12.	Liguori C, Nuccetelli M, Izzi F, et al. Rapid eye movement sleep disruption and sleep fragmentation are associated with increased orexin-A cerebrospinal-fluid levels in mild cognitive impairment due to Alzheimer's disease. *Neurobiol Aging.* 2016;40:120-126.

13.	Beaulieu-Bonneau S, Hudon C. Sleep disturbances in older adults with mild cognitive impairment. *Int Psychogeriatr.* 2009;21(4):654-666.

14.	Cross N, Terpening Z, Rogers NL, et al. Napping in older people 'at risk' of dementia: relationships with depression, cognition, medical burden and sleep quality. *J Sleep Res.* 2015;24(5):494-502.

15.	da Silva RA. Sleep disturbances and mild cognitive impairment: A review. *Sleep Sci.* 2015;8(1):36-41.

16.	Guarnieri B, Adorni F, Musicco M, et al. Prevalence of sleep disturbances in mild cognitive impairment and dementing disorders: a multicenter Italian clinical cross-sectional study on 431 patients. *Dement Geriatr Cogn Disord.* 2012;33(1):50-58.

17.	Lambert JC, Ibrahim-Verbaas CA, Harold D, et al. Meta-analysis of 74,046 individuals identifies 11 new susceptibility loci for Alzheimer's disease. *Nat Genet.* 2013;45(12):1452-1458.

18.	Zetterberg H, Alexander DM, Spandidos DA, Blennow K. Additional evidence for antagonistic pleiotropic effects of APOE. *Alzheimers Dement.* 2009;5(1):75.

19.	Barnes J, Dickerson BC, Frost C, Jiskoot LC, Wolk D, van der Flier WM. Alzheimer's disease first symptoms are age dependent: Evidence from the NACC dataset. *Alzheimers Dement.* 2015;11(11):1349-1357.

20.	Mistridis P, Krumm S, Monsch AU, Berres M, Taylor KI. The 12 Years Preceding Mild Cognitive Impairment Due to Alzheimer's Disease: The Temporal Emergence of Cognitive Decline. *J Alzheimers Dis.* 2015;48(4):1095-1107.

21.	Hewitt J, Walters M, Padmanabhan S, Dawson J. Cohort profile of the UK Biobank: diagnosis and characteristics of cerebrovascular disease. *BMJ Open.* 2016;6(3):e009161.

22.	Fry A, Littlejohns TJ, Sudlow C, et al. Comparison of Sociodemographic and Health-Related Characteristics of UK Biobank Participants With Those of the General Population. *Am J Epidemiol.* 2017;186(9):1026-1034.

23.	Krokstad S, Langhammer A, Hveem K, et al. Cohort Profile: the HUNT Study, Norway. *Int J Epidemiol.* 2013;42(4):968-977.
